# Pelage Variation and Morphometrics of Closely Related *Callithrix* Marmoset Species and Their Hybrids

**DOI:** 10.1101/2023.01.01.522211

**Authors:** Joanna Malukiewicz, Kerryn Warren, Vanner Boere, Illaira LC Bandeira, Nelson HA Curi, Fabio T das Dores, Lilian S Fitorra, Haroldo R Furuya, Claudia S Igayara, Liliane Milanelo, Silvia B Moreira, Camila V Molina, Marcello S Nardi, Patricia A Nicola, Marcelo Passamani, Valeria S Pedro, Luiz CM Pereira, Bruno Petri, Alcides Pissinatti, Adriana Alves Quirino, Jeffrey Rogers, Carlos R Ruiz-Miranda, Daniel L Silva, Ita O Silva, Monique OM Silva, Juliana L Summa, Ticiana Zwarg, Rebecca R Ackermann

## Abstract

**Background:** Hybrids are expected to show greater phenotypic variation than their parental species, yet how hybrid phenotype expression varies with genetic distances in closely-related parental species remains surprisingly understudied. Here we study pelage and morphometric trait variation in anthropogenic hybrids vailable at the end of the article between four species of Brazilian *Callithrix* marmosets, a relatively recent primate radiation. Marmoset species are distinguishable by pelage phenotype and morphological specializations for eating tree exudates. Here, we (1) describe qualitative phenotypic pelage differences between parental species and hybrids; (2) test whether significant quantitative differences exist between parental and hybrid morphometric phenotypes; and (3) determine which hybrid morphometic traits show heterosis, dysgenesis, trangression, or intermediacy relative to the parental trait. For morphometric traits, we investigated both cranial and post-cranial traits, particularly as most hybrid morphological studies focus on the former instead of the latter. Finally, we estimate mitogenomic distances between marmoset species from previously published data.

**Results:** Marmoset hybrid facial and overall body pelage variation reflected novel combinations of coloration and patterns present in parental species. In morphometric traits, *C. jacchus* and *C. penicillata* were the most similar to each other, while *C. aurita* was the most distinct, and *C. geoffroyi* trait measures fell between these other species. Only three traits in *C. jacchus* x *C. penicillata* hybrids showed heterosis. We observed heterosis and dysgenesis in several traits of *C. penicillata* x *C. geoffroyi* hybrids. Transgressive segregation was observed in hybrids of *C. aurita* and the other species. These hybrids were also *C. aurita*-like for a number of traits. Genetic distance was closest between *C. jacchus* and *C. penicillata* and farthest between *C. aurita* and the other species.

**Conclusion:** We attributed significant morphometric differences between marmoset species to variable levels of morphological specialization for exudivory in these species. Our results suggest that intermediate or parental species-like hybrid traits relative to the parental trait values are more likely in crosses between species with relatively lesser genetic distance. More extreme phenotypic variation is more likely in parental species with greater genetic distance, with transgressive traits appearing in hybrids of the most genetically distant parental species. We further suggest that less developmental disturbances can be expected in hybrids of more recently diverged parental species, and that future studies of hybrid phenotypic variation should investigate selective pressures on *Callithrix* cranial and post-cranial morphological traits.

## Background

Hybridization occurs under both natural and anthropogenic contexts, with the former occurring in about 10% of animal species [1], and with the latter increasing between previously isolated populations [2, 3, 4]. Our understanding of the genomic consequences of animal hybridization has grown considerably (e.g.[4, 5, 6, 7]), and the range of hybridization outcomes include but are not limited to hybrid speciation (the origin of a new species via hybridization between two distinct species), genetic swamping (maladaptive gene flow from central populations into peripheral populations [8]), adaptive introgression (the incorporation of a foreign genetic variant via hybridization from a donor pool that leads to an increase of the fitness of the recipient pool [9]), or extinction (the termination of a genetic lineage) [5, 6, 7]. Hybridization also impacts morphological traits [10, 11, 12]. Studies of hybrid morphology to date have largely focused on craniofacial features, but we still possess knowledge gaps in how hybridization manifests itself in post-cranial anatomy [12]. Given the importance of post-cranial morphology in locomotion and reproduction and that different selective forces likely act on post-cranial and cranial morphology [13], hybrids may express cranial traits differently than post-cranial traits. Animal hybrid morphology studies also feature a single pair of parental species and the resulting hybrids (e.g. [10, 14, 15, 16, 17, 18, 19]), but there is also interest in understanding how the hybrid phenotype varies with the genetic distances between closely-related parental species [10, 20].

Hybrids are expected to show a more variable array of morphological phenotypes than their parental species [10, 21]. Hybrids can resemble one of their parental species, either in terms of a single trait or as a whole, can be heterotic or dysgenetic relative to the parents (measured as positive or negative deviation from a mid-point value), or can display transgressive traits (i.e. outside of the range of parental variation)[10, 21, 22]. The cumulative effects of gene interactions (dominance and epistasis), parental species temporal divergence, and allele frequency differences between parental species are all thought to underlie morphological phenotypic variation in hybrids [21]. Intermediate traits are explained by a standard polygenic model with additive effects, which is expected for species with small allele frequency differences [10, 21]. However, isolated parental populations with different fixed alleles are expected to produce heterotic hybrids [10, 21]. Dysgenesis is predicted for more distantly related taxa and represents a breakdown of ‘coadapted gene complexes’ between the parental species [10, 21]. Transgressive traits seem to be related to complementary gene action of antagonistic quantitative trait loci [23, 24]. Thus, the frequency of heterosis, dysgenesis, and trangressive segregation in hybrid populations should increase with greater genetic distance between parental species, as longer divergence times allow for more fixation of complementary alleles in parental populations.

As pointed out by Ackermann [10], a lingering question about the evolutionary importance of hybrid phenotypic expression is “to what extent might differences in the expression of hybrid traits exist due to degree of temporal divergence?” One key study which looked at the phenotypic effects of hybridization in pairs of parental species within a wide range of genetic distance was conducted experimentally on cichlid fish [20], and there was a particular interest in transgressive traits in this work. In F1 hybrids, the relationship between the frequency of transgressive segregation and level of parental species genetic difference had a concave shape while in F2 hybrids the amount of hybrid transgression increased linearly with parental species genetic distance [20]. However beyond such work, hybrid expression of morphological traits across interbreeding species with variable genetic difference, particularly in non-experimental animal populations, remains understudied.

Primates are one animal group where hybridization is estimated to occur among 7–10% of species [25], and the recent radiation of Brazilian *Callithrix* marmoset makes an excellent model for characterizing hybridization effects between closely-related species with variable degrees of temporal divergence. The two phylogenetic subgroups that compose the *Callithrix* genus, the “*aurita*” group (*C. aurita* and *C. flaviceps*) and the “*jacchus*” group (*C. kuhlii*, *C. geoffroyi*, *C. jacchus*, *C. penicillata*), diverged about 3.5 million years ago (Ma) [26]. Within the *jacchus* group, *C. jacchus* and *C. penicillata* are the most recently diverged at 0.51 Ma, followed by *C. kuhlii* at 0.82 Ma, and *C. geoffroyi* at 1.18 Ma [27]. *Callithrix* species are distinguishable from each other based on level of morphological specialization for eating tree gums and exudates (ie. exudivory), facial and overall body pelage patterns and coloration, and peri-auricular ear-tuft shape and color [27]. Limited *Callithrix* hybridization already occurs naturally between certain pairs of *Callithrix* species like *C. jacchus* and *C. penicillata* under secondary contact at species range boundaries, however the illegal pet trade has dramatically increased anthropogenic *Callithrix* hybridization relatively to natural conditions [26, 27, 28].

Thus far, most studies of hybrid *Callithrix* phenotypes are based on qualitative descriptions of pelage differences between hybrids and their parental species [29, 30, 31, 32, 33, 34]. Only Fuzessy et al. [35] and Cezar et al. [36] have tested theoretical expectations of hybrid phenotypic diversity in *C. geoffroyi* x *C. penicillata* and *C. jacchus* x *C. penicillata* hybrids, respectively. Here, we build upon these previous studies by examining cranial and post-cranial metric variation among four marmoset species (*C. aurita*, *C. jacchus*, *C. geoffroyi*, *C. penicillata*) along with their hybrids in individuals sampled in the wild or in captivity. Our study represents the largest marmoset morphological sampling to date in terms of hybrid sample number and types of hybrids.

Our main study aims are to: (1) describe qualitative pelage phenotypic differences between parental species and hybrids; (2) test whether significant quantitative differences exist between parental and hybrid marmoset phenotypes; (3) quantify whether and how hybrid phenotypic variation differs relative to parental species (i.e., intermediate, heterotic, dysgenic, or transgressive); and (4) investigate how aims 2 and 3 vary with differential parental species’ genetic distance, which we use as a proxy for temporal divergence. We estimated genetic distances between marmoset species from previously published mitogenomic data that include a subset of our samples [26]. Based on these aims, our first hypotheses is that the highest occurrence of intermediate morphological traits exists between *C. jacchus* and *C. penicillata* hybrids, as their parental species as the two most recently diverged within *Callithrix*. Given longer divergence times between *jacchus* and *aurita* group species than between *jacchus* group species, we hypothesize that dysgeneic and/or transgressive traits appear more frequently in hybrids of the former than in the latter set of species.

## Methods

### Sampling

Our samples consisted of 209 adult individuals (Table 1 and Supplementary Table S1) from four *Callithrix* species (*C. aurita, C. geofforyi, C. jacchus, C. penicillata*) as well as several hybrid types (*C. aurita* x *Callithrix* sp., *C. penicillata* x *C. geoffroyi*, *C. penicillata* x *C. jacchus*, *Callithrix* sp. x *Callithrix* sp). Following Yamamoto [37] observations of dental characteristics and genitalia growth in marmosets, animals between 5 and 10 months old were classified as juveniles, while those older than 11 months were considered adults. We excluded all non-adult individuals from the phenotypic and morphological analyses described below.

**Table 1.** Marmoset sample size by taxon. “N” represents the number of individuals sampled for each given taxon.

| Taxon | N |
| --- | --- |
| *C. aurita* | 27 |
| *C. aurita* × *Callithrix* sp. | 9 |
| *Callithrix* sp. × *Callithrix* sp. | 2 |
| *C. geoffroyi* | 14 |
| *C. jacchus* | 30 |
| *C. penicillata* | 55 |
| *C. penicillata* × *C. geoffroyi* | 18 |
| *C. penicillata* × *C. jacchus* | 54 |

Marmosets were sampled between 2015 and 2019 as follows: (1) wild marmosets in Bahia, Espírito Santo, Minas Gerais, Rio de Janeiro, Pernambuco, and São Paulo states; (2) captive-born, wild-caught, and confiscated marmosets housed at the Guarulhos Municipal Zoo, Guarulhos, São Paulo, CEMAFAUNA (Centro de Manejo de Fauna da Caatinga), Petrolina, Pernambuco, CPRJ (Centro do Primatologia do Rio de Janeiro), Guapimirim, Rio de Janeiro, Parque Ecol^ogico do Tiet^e (PET), São Paulo, SP, and Divisão Técnica de Medicina Veterińaria e Manejo da Fauna Silvestre (DEPAVE-3), São Paulo, SP; (3) a wild group from Natividade, Rio de Janeiro that was caught and housed at CPRJ; and (4) a wild group from Ilha D’Agua, Rio de Janeiro, RJ housed at SERCAS (Setor de Etologia aplicada à Reintrodução e Conservação de Animais Silvestres), Campos dos Goytacazes, RJ. Marmoset capture methodology has been described elsewhere [34]. All individuals were allowed to recover after sample collection, and wild marmosets were released at their original point of capture.

### Phenotyping of <u>Callithrix</u> Species and Hybrids (Aim 1)

Using the approach developed in Fuzessy et al. [35], marmoset facial markings and pelage characteristics were used to phenotypically differentiate between species and hybrids (Supplementary Figure S1). Defining facial and pelage characteristics from each species and hybrid type were based on published descriptions [30, 35, 34, 38, 28] and personal observations by JM and CSI. Phenotypes of hybrids classified as *C. aurita* hybrids suggest that these individuals possess ancestry from *C. aurita* and at least one species from the *jacchus* group [28, 38]. Previous phylogenetic analysis of mitogenomic haplotypes assigned to a subset of *C. aurita* hybrids used in our sample also support *C. aurita* x *jacchus* group ancestry in these individuals (BJT024/*C. aurita* mitogenome, BJT025/*C. jacchus* mitogenome, BJT026/*C. penicillata* mitogenome, BJT027/*C. geoffroyi* mitogenome, BJT115/*C. aurita* mitogenome) [27]. Two hybrids were not able to be classified at the species level due to ambigious phenotypes, and were therefore classified as *Callithrix* sp. x *Callithrix* sp. hybrids. The only exception was hybrid BJT070 for which previous mitogenomic phylogenetic analysis determined *C. geoffroyi* to be one of the parental species [27].

**Figure 1.**
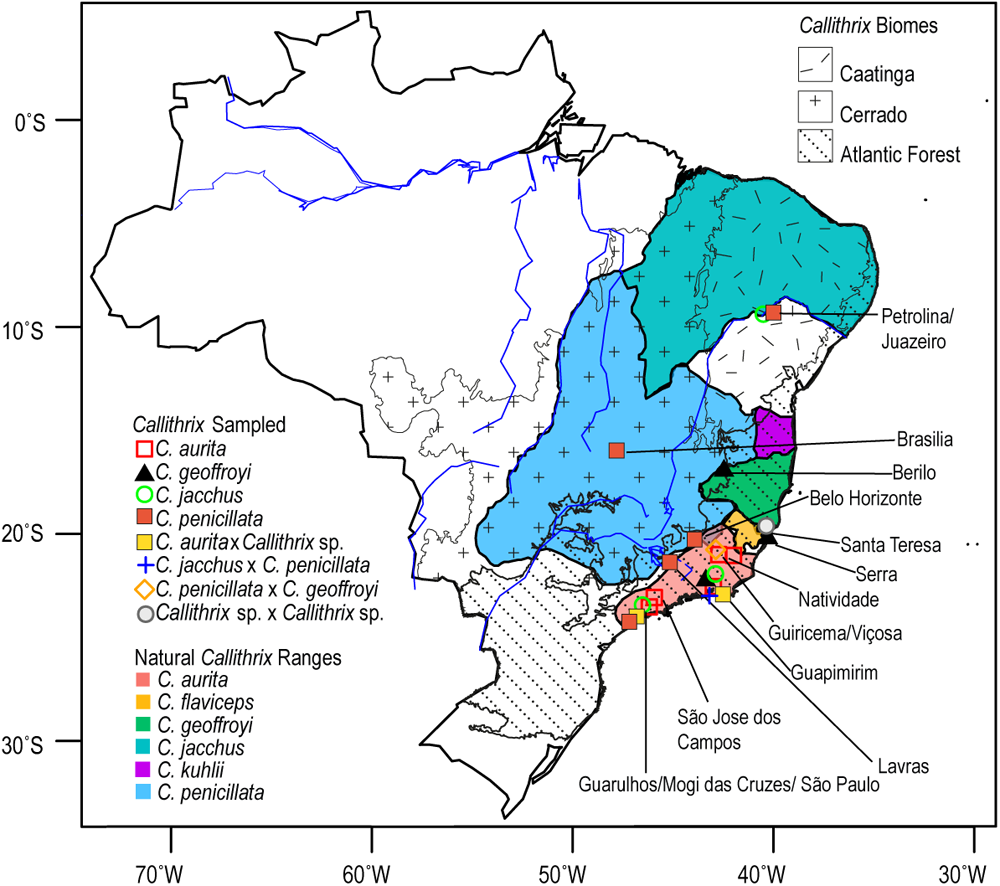
Marmoset sampling locations. Sampling locations are indicated by di_erent color symbols, and the approximate natural distribution of Callithrix species in Brazil are identi_ed by di_erent colors. The distribution maps are based on 2012 IUCN Red List Spatial Data (http://www.iucnredlist.org/technical-documents/spatial-data). The locations of the three biomes where Callithrix occur naturally, the Caatinga, Cerrado, and Atlantic Forest, are also indicated with di_erent patterning

### Quantitative Testing of <u>Callithrix</u> Morphometric Trait Differences Between <u>Callithrix</u> Species and Hybrids (Aim 2)

Sampled adults were measured with a tape measure and digital calipers and weighed while under anesthesia, following methods described by Nagorsen and Peterson [39]. Metric data are represented by one measure of body weight (WEIGHT) taken in grams (g), and 12 linear distances (Supplementary Table S1). Linear distances measured in centimeters (cm) were tail length (TAIL), humeral length (HUMERUS), forearm length (FOREARM), body length (BODY), femur length (FEMUR), tibia length (TIBIA). Linear distances measured in millimeters (mm) were maximal intercranial distance (IC), fronto-occipital distance (FO), widest distance between zygomatic arches (ZYG), distance between mandible angles (JAW), wrist-longest claw (HAND), and calcaneus-longest claw (FOOT). For HAND, HUMERUS, FOREARM, FEMUR, TIBIA, and FOOT measures, we measured both left and right sides on sampled individuals, and then took the bilateral average of each measurement for further analyses.

All analyses described below were carried out in R [40] and code is available in Supplementary File “Morphometricsv3 code.Rmd.” To first check for normality of the data, we produced normal quantile-quantile (QQ) plots for all variables. For each variable most points fall approximately along the reference line (Supplementary Figure S2). We also inspected stem-and-leaf plots for each variable (see Results). Although some variables indicated slight deviation from normality based on these plots, the parametric statistical tests described below are fairly robust to such violation, so we left the measured traits uncorrected [41].

To test for any confounding effects from sexual dimorphism in our data, we conducted a series of parametric multivariate analysis of variance (MANOVA). We first used MANOVA to test for an interaction between sex and taxon for all 13 morphological traits, which was not statistically significant (p-value=0.9665). Grouping all 13 traits by sex indicated that these variables do not differ significantly between males and females (p-value=0.74). On the other hand, grouping all 13 traits by taxon in the MANOVA test indicated a statistically significant effect of taxon (parametric MANOVA F(91, 910) = 2.7957, p*<*0.01) these MANOVA tests, we do not expect there to be any confounding effects from sexual dimorphism on the thirteen morphological traits in our data set.

Following these tests, each of the 13 measurements was analyzed individually using ANOVA to test for differences between all taxa. Prior to running each ANOVA test, we checked for homogeneity of variances by Levene’s test for each variable among taxa. Levene’s test indicated that the BODY, IC, FO, FOREARM, FEMUR, TIBIA, and FOOT traits had homogeneity of variance with p-value *>*0.05. All other traits produced significant p-values (*<*0.05) for Levene’s test. As not all traits showed homogeneity of variance (see Results), we conducted one-way Welch’s ANOVAs, which were followed up by Games-Howell post-hoc tests to perform multiple pairwise comparisons between groups. Prior to conducting univariate ANOVA tests, we generated normality QQ plots for each respective trait (Figure S2). The Games-Howell test was carried out with the Rstatix [42] R package and p-values were adjusted for multiple comparisons using the Tukey method.

### Quantitative Testing for Intermediacy, Heterosis, Dysgenesis, and Transgressive Segregation of Morphometric Traits in <u>Callithrix</u> Hybrids (Aim 3)

For *C. jacchus* x *C. penicillata*, *C. penicillata* x *C. geoffroyi*, and *C. aurita* hybrids, we compared hybrids and parental species to determine if any traits showed evidence of heterosis, dysgenesis, or transgressive segregation. For *C. aurita* hybrids, all possible combinations of *C. aurita* and *jacchus* group species from our samples were used as putative parental species as it was not possible to determine the exact parental species of *C. aurita* hybrids. Other hybrid types were excluded from these tests due to relatively small sample numbers. First, we calculated the mid-point values (MPVs) for each possible parental pair of species for all 13 traits. MPVs for each trait were calculated multiplying the sum of parental species means for each trait by 0.5. We then compared trait means of each hybrid group against their respective MPVs using one-sample t-tests. Mean hybrid trait values that fell in between parental trait means and were not statistically significantly different from the MPVs were considered intermediate. Mean hybrid trait values that considered parental-like for a given parental species when the hybrid trait mean was closer to mean trait values of a given parental species and were not statistically significantly different from the MPVs. Mean hybrid trait values that were significantly larger than the MPVs were considered heterotic. Mean hybrid trait values significantly smaller than the MPVs were considered dysgenic. Following this, Welch’s two sample t-tests, which account for unbalanced size and lack of variance homogeneity among samples, were conducted between trait means of hybrids and each parental species. A trait was considered transgressive if the hybrid mean was larger than both parental means, and all hybrid-parental species Welch’s t-tests were statistically significant.

A principal components analysis (PCA) was also performed on the data in order to visualize differences among the species and hybrids. This technique reduces the dimensionality of a data set producing a smaller number of uncorrelated variables that nonetheless retain all of the original size and shape information. Separate PCAs were conducted for *C. jacchus* x *C. penicillata*, *C. penicillata* x *C. geoffroyi*, and *C. aurita* hybrids. For *C. aurita* hybrids, as described above, all possible combinations of *C. aurita* and *jacchus* group species from our samples were used as putative parental species

### Genetic Distance between <u>Callithrix</u> Species (Aim 4)

To determine mean pairwise genetic distances between *C. aurita*, *C. jacchus*, *C. penicillata*, and *C. geoffroyi*, we used previously published mitogenomic sequences [26], which included a subset of marmosets used in this current study. Samples and mitogenomic Genbank accession numbers are listed in Supplementary Table S2. Mitogenomic haplotypes were grouped by species and mean genetic distances between these groups were calculated with MEGA11 [43, 44]. We used the “Compute Between Group Mean Distance” option with default settings of the Maximum Composite Likelihood model, transitions and transversions substitutions included, uniform rates among sites, same (homogeneous) patterns among lineages, and pairwise deletion as gaps/missing data treatment.

## Results

### Descriptions of <u>Callithrix</u> Phenotypes (Aim 1)

#### <u>Callithrix</u> Species Phenotypes

Examples of the *C. aurita* phenotype are shown in Fig. 2A and summarized in Supplementary Table S3. The frontal half vertex of *C. aurita* varies between beige, orange, and black and the back half of the vertex varies from orange to black. The menton region has yellowish to orange pelage, while the orbital region contains a mix of yellowish and peachy pelage. The *C. aurita* ear tufts frame the facial region but the tuft hair is not as full or dense in volume as that of *C. jacchus*; the ear tufts may be yellow or orange. The pelage of the *C. aurita* facial lateral sides is black. The forehead, nasal, and infraorbital regions have beige to light orange pelage. Pelage on the back does not form a pattern of obvious striae, but proximally there is a mixture of orange banded patches (the orange is more intense than that of *C. jacchus* and *C. penicillata*) among black pelage. The orange coloration of the back is less intense moving proximal to distal, and becomes predominately black towards the tail base. The proximal region of the neck has black hair, but the distal region has pelage that follows the pattern described for the back. The belly region has black pelage with some slightly orange tips at the distal part of the hairs. The proximal regions of the arms and legs have black pelage with some with orange tips. The distal base of the arm has also black hair with orange tips that is more evident than in the distal part of the legs. The tail pelage has a black, grey, and orange striated pattern.

**Figure 2.**
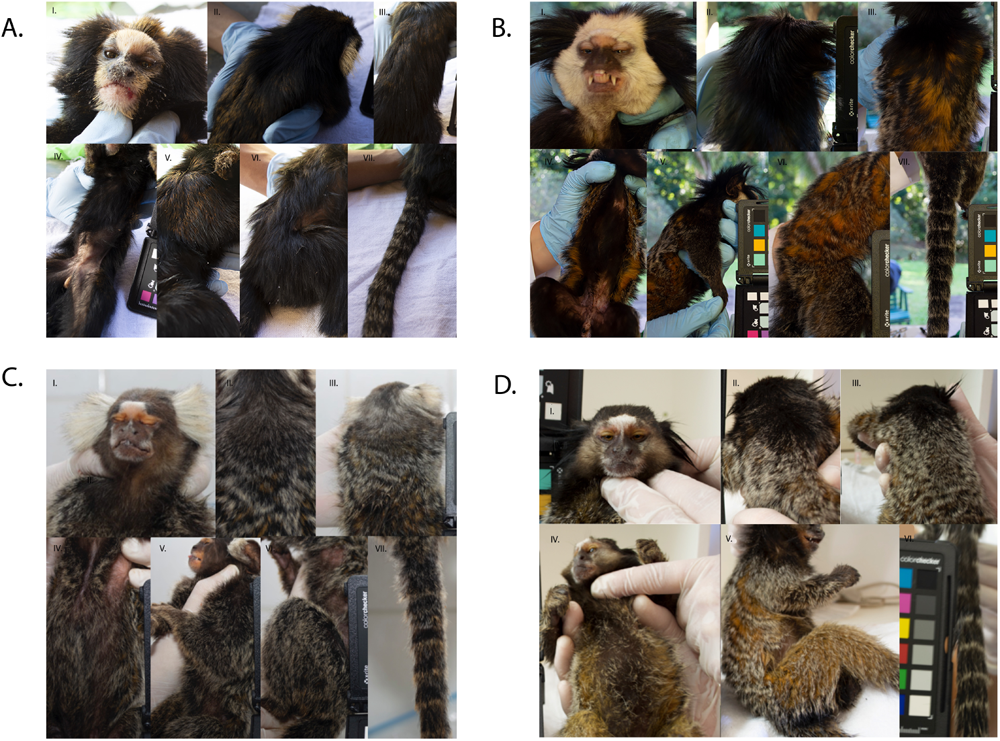
Phenotypes of four Callithrix species. Partition A shows the C. aurita face and ear tufts (I), neck and upper back (II), full back (III), belly (IV), arm (V), leg (VI), and tail (VII). Partition B shows the C. geo_royi face and ear tufts (I), neck (II), full back (III), belly (IV), arm (V), leg (VI), and tail (VII). Partition C shows the C. jacchus face and ear tufts (I), neck and upper back (II), full back (III), belly (IV), arm (V), leg (VI), and tail (VII). Partition D shows the C. penicillata face and ear tufts (I), neck and upper back (II), back (III) belly (IV), arm and leg (V), and tail (VI).

The *C. geoffroyi* phenotype is shown in Fig. 2B and summarized in Supplementary Table S3. The front half of the vertex of *C. geoffroyi* is fully white while the back half of the vertex and proximal portion of the head is black. The orbital region is peachy, but the forehead and most of the face around the orbital, nasal, and infraorbital regions are also white. The pelage of the menton region can be white or beige combined with darker hairs. The *C. geoffroyi* ear tuft pelage is very dense as in *C. jacchus*, and similar in volume, but the ear tuft hair is black. Tuft hairs closer to the top of the head are shorter and tuft hairs closer to the neck are longer. The neck pelage is black, and the back region has striations which can be either black and orange or black and grey. Portions of orange coloration in the pelage of the back are obvious and prominent. The proximal portions of the arms and legs are black and can be speckled with a whitish-grey coloration with overall darker coloring on the outer parts in the arms and legs. Tail pelage has a black, grey and orange striated pattern.

The *C. jacchus* phenotype is shown in Fig. 2C and summarized in Supplementary Table S3. *Callithrix jacchus* pelage of the front half of the vertex is dominated by grey tips of hair, but can also have beige or brown tones. The back portion of the vertex is brown with tips of grey hair. The pelage of the menton region is grey. The facial orbital region is more peachy and buff colored than in *C. penicillata*. The *C. jacchus* tufts are periauricular, white and the hair is highly voluminous. Tips of the *C. jacchus* tuft hairs may have some black tones. The pelage on the lateral sides of face ranges from dark brown to a little orange with some hairs that may have greyish tips. A white ‘star’ is present and prominent on the forehead of *C. jacchus*. The upper neck region has dark brown coloration, while the lower neck region transitions towards aguti coloration. Striations present with black and whitish-grey topcoat pelage and an orange colored pelage undercoat define the back region. The arms have black to dark brown pelage and tips of pelage hairs are grey to light orange or orange. The legs follow the striation pattern of the back region. Tail pelage has black, grey and orange striated pattern.

The *C. penicillata* phenotype is shown in Fig. 2D and summarized in Supplementary Table S3. The front and back halves of the vertex pelage are dark brown to black. The pelage of the menton region is whitish-grey, while the facial orbital region pelage is creme-buffy colored. The ear tufts are preauricular and this region has thin, downward facing, relatively long black pelage. There is a prominent white ‘star’ present on the *C. penicillata* forehead and the pelage on the lateral sides of face is whitish-grey to dark brown. The upper and lower neck pelage has dark brown and black coloration, with occasional presence of specks of whitish-grey. Striations on the back combine a whitish-grey/black pelage topcoat with an orange pelage undercoat. Light-orange to orange and black pelage is present in the central belly region of *C. penicillata*. The proximal region of the arms is predominantly whitish-grey, and the proximal region of the legs follows the striation pattern of back region. Tail pelage has black and whitish-grey striations.

#### <u>Callithrix</u> Hybrid Phenotypes

Examples of anthropogenic *C. jacchus* x *C. penicillata* hybrid phenotypes from southeastern Brazil are shown in Fig. 3A and summarized in Supplementary Table S3. The front half of the vertex of *C. jacchus* x *C. penicillata* hybrids is composed of grey and black hair of varying intensities, while the back half of the vertex may range from black to greyish and/or orange pelage. The menton region pelage is grey. Pelage of the orbital region has variable shades of orange, and may even be pink. The ear tuft pelage of *C. jacchus* x *C. penicillata* hybrids is usually less voluminous than in *C. jacchus* but more so than in *C. penicillata*. Hybrid ear-tuft coloration ranges from black with grey tips to grey with some black hair. These hybrids have a white ‘star’ present on the forehead, as also possessed by parental *C. jacchus* and *C. penicillata*, but the hybrid star mark varies in size. The lateral sides of the face of hybrids have pelage of greyish coloration with some black and orange hairs. Coloration of neck pelage may be black, grey, and/or orange. Hybrid back pelage has striations interspersed with orange, black, and grey coloration. The striation patterns may not be as uniform as in parental species. The intensity of orange back coloration varies among hybrid individuals. The belly pelage varies in intensity from black to orange, but these two colors are striated. Pelage on the proximal region of the legs follows the pattern of the back region. The proximal regions of the arms have black to dark brown fur with grey tips. The tail pelage has black, grey and orange striated pattern, varying in color intensity.

**Figure 3.**
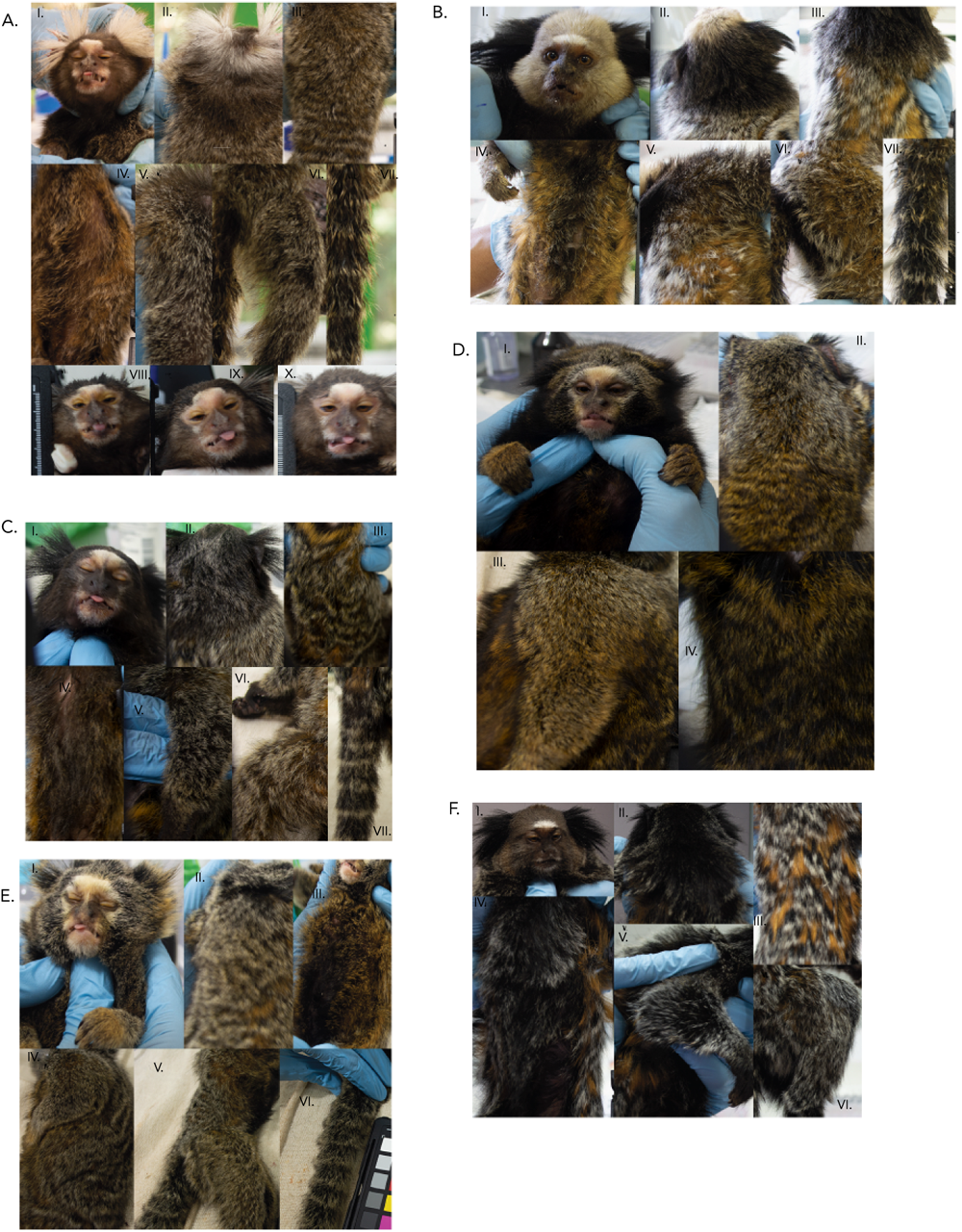
Phenotypes of Callithrix hybrids. Partition A shows examples of C. jacchus x C. penicillata hybrid face and ear tufts (I), neck and upper back (II), back (III), belly (IV), arm (V), leg (VI), tail (VII), and further facial variation (VIII-X). Partition B shows examples of C. penicillata x C. geo_royi hybrid face and ear tufts (I), neck (II), back (III), belly (IV), arm in upper right of photograph (V), leg (VI), and tail (VII). Partition C shows an example of a C. aurita hybrid phenotype for face and ear tufts (I), neck and upper back (II), full back (III), belly (IV), arm (V), leg (VI), and tail (VII). Partition D shows an example of another C. aurita hybrid phenotype for face and ear tufts (I), neck and upper back (II), arm (III), and belly (IV). Part E shows an example of another C. aurita hybrid phenotype for face and ear tufts (I), neck and upper back (II), belly (III), arm in upper portion of photograph (IV), leg in lower portion of photograph (V), and tail (VI). Partition F shows an example of a C. geo_royi x Callithrix sp. hybrid phenotype for face and ear tufts (I), neck and upper back (II), back (III), belly(IV), arm (V), and leg (VI).

Examples of anthropogenic *C. geoffroyi* x *C. penicillata* hybrid phenotypes from Viçosa, Minas Gerais are shown in Fig. 3B and summarized in Supplementary Table S3. For these hybrids, the pelage of the front half of the vertex, back half of the vertex, and lateral sides of the face varies in intensity from white to grey. Pelage of the upper neck of the hybrids varies from white to dark grey. In the lower neck part, the hair can be black and may have grey tips. In the facial menton region of hybrids, pelage follows the pattern of lateral sides of the face. In the facial orbital region, hybrids have pelage that is slightly orange or peachy. The hybrid ear tuft pelage color is black but the volume of tufts varies between that of the parental species. The white forehead mark of *C. penicillata* is present in these hybrids but varies in intensity between individual hybrids. The pelage of the back region possesses patterns of black, grey and orange streaks, as seen in the parental species. Black hairs are found in the central part of the belly, but the hairs are intense orange in the outer parts of the belly. The proximal portion of the legs follows the pelage pattern of the back, and the proximal portion of the arms has black hairs with some grey tips. The tail pelage shows a black, grey and orange striated pattern.

Examples of anthropogenic *C. aurita* x *Callithrix* sp. hybrid phenotypes are shown in Fig. 3C-E and summarized in Supplementary Table S3. *Callithrix aurita* x *Callithrix* sp. hybrids have a front vertex half with black and grey hairs that have orange tips. In the back half of the vertex, the pelage coloration contains black hair with grey tips, with variation in the intensity of the grey. The vertex of some hybrid individuals will have patches of whitish-grey and grey mixed in with the darker black pelage hairs. This pattern also occurs in the neck region. The menton region pelage is whitish-grey, and the orbital region pelage may be peachy as in *C. jacchus* and *C. penicillata*, or yellowish like *C. aurita*. Hybrid ear tuft hair volume may be sparse like *C. aurita* and *C. penicillata* or very dense like *C. jacchus*, varying in the amount of black, grey, and orange hair at the hair tips. Some hybrids possess a white star on the forehead. Others will have a *C. aurita*-like pattern where the forehead, orbital, nasal, infraorbital, and menton facial regions have beige to light orange hairs. The lateral sides of the face have black to dark brown hair that may or may not have grey tips.

Unlike *C. aurita*, *C. aurita* x *Callithrix* sp. hybrids show back striation patterns that are similar to that of *C. penicillata* and *C. jacchus*. The striations may contain a mixture of black, grey and orange patterns or black and whitish-grey streaks. In *C. aurita* x *Callithrix* sp. hybrids, the orange color of back pelage tends to be more intense than in *C. aurita*, and greys of the back pelage are more yellowish or orange instead of whitish than in *C. penicillata* and *C. jacchus*. Belly coloration is highly variable between hybrids. The proximal region of legs follows the pattern of the back. The proximal portion of the arm has black fur with grey to orange tips. The hybrid tail pelage has a black and grey striated pattern and there may be orange coloration at hair tips. The hands of these hybrids tend to have an orange or yellow tone, similar to *C. aurita*.

An example of a *Callithrix* sp. x *Callithrix* sp. hybrid phenotype from Santa Teresa, Espírito Santo is shown in Fig. 3F and summarized in Supplementary Table S3. For this hybrid, the front of vertex pelage is yellowish with a mix of grey and black speckles, and the back of the vertex pelage is black with greyish speckles. Pelage of the facial menton region is dark. The facial orbital region pelage is black towards the eyes and peachy on the outer regions. A white forehead star is present in these hybrids. The ear tuft pelage is very dense as in *C. geoffroyi*. Hybrid ear tufts are black, and hairs closer to top of the head are shorter and hairs closer to the neck are longer. The upper neck region has black hair, while the lower neck portion has greyish tips. Pelage in the back has striations that are black*\*orange and black*\*grey. The orange coloration is very obvious and prominent in the hairs of the back pelage. The belly pelage contains striations of black and grey. The proximal leg portion is black and the proximal arm region has whitish-grey hairs. The individual pictured in Figure 3F likely possesses ancestry from *C. penicillata* or *C. jacchus* given the forehead star, as well as previously confirmed *C. geoffroyi* ancestry. However, this phenotype is distinct from that described for *C. penicillata* x *C. geoffroyi* hybrids described above.

### Quantitative Differences between Parental and Hybrid Callithrix Morphometric Traits (Aim 2)

Univariate Welch’s ANOVA tests (Supplementary Table S4) indicate significant differences between mean trait values among *Callithrix* taxa. Among species, we consistently see significant differences between *C. aurita* and *C. jacchus* and *C. penicillata*, respectively across most mean trait values (Supplementary Table S5 and Fig. 4). For most traits, *Callithrix aurita* was the largest *Callithrix* species, as it tended to have the highest trait median and mean values across all taxa (Table 2 and Fig. 4). Only in the HAND trait did post-hoc tests fail to find significant differences in pairwise comparisons among species (Supplementary Table S5 and Fig. 4). Within the *jacchus* group, *C. geoffroyi* tended to be the largest for most traits (Table 2 and Fig. 4). Additionally, *C. geoffroyi* was significantly different for a larger number of traits when compared with *C. jacchus* than with *C. penicillata* (Supplementary Table S5 and Fig. 4). The respective FEMUR, TIBIA, and HUMERUS means of *C. geoffroyi* were significantly different from that of both *C. jacchus* and *C. penicillata* (Supplementary Table S5 and Fig. 4). There were no significant differences between

**Figure 4.**
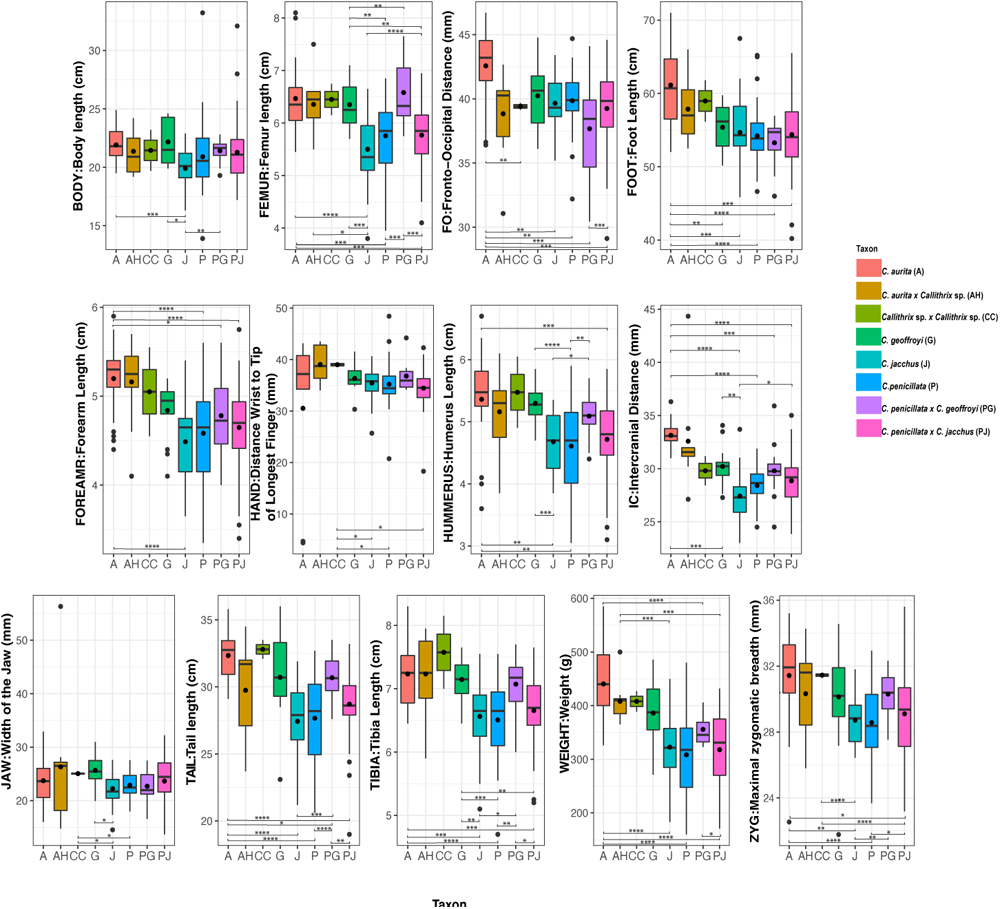
Stem and leaf box plots for 13 morphological traits in four Callithrix species and their hybrids. The x-axis of each plot represents Callithrix taxon categories and the y-axis of each line represents values of trait measurements. Boxes represent the respective interquartile ranges of 13 Callithrix morphological trails. The bottom box lines represent 25th percentiles, the mid-lines of boxes represents 50th percentiles/medians, and top box lines represents 75th percentiles. Dots inside of box represent respective trait means, and dots outside of respective boxes represent trait outliers. Bottom whiskers of each box represent the variability of minimum trait values relative to the interquartile range and the top whiskers of each box represent maximum trait values relative to the interquartile range. Signi_cant p-values for taxon di_erences from Supplementary Table S5 Games-Howell post-hoc pairwise tests results following Welch’s ANOVA are represented by as “*” for p-value<0.05, as “**” for p-value<0.01, and as “***” for p-value<0.001. Taxon abbreviations as well as the along the x-axis in each plot and the _gure legend are as follows: AC. aurita, G-C. geo_royi, J-C. jacchus, P-C. penicillata, AH-C. aurita x Callithrix sp. hybrid; CC-Callithrix sp. x Callithrix sp. hybrid; PG-C. geo_royi x C. penicillata hybrid; PJ-C. penicillata x C. jacchus hybrid. “G” stands for grams, “cm” stands for centimeters, and “mm” stands for millimeters.

**Table 2.** Summary of species means, standard deviations (SD), and sample numbers (N) of thirteen *Callithrix* morphological traits. “Cm” refers to centimeters, “g” to grams, “mm” to millimeters, “A” to *C. aurita*, “G” to *C. geoffroyi*, “P” to *C. penicillata*, and J to *C. jacchus*.

| Trait | *C. aurita* (A) |  |  | *C. geoffroyi* (G) |  |  | *C. jacchus* (J) |  |  | *C. penicillata* (P) |  |  |
| --- | --- | --- | --- | --- | --- | --- | --- | --- | --- | --- | --- | --- |
|  | N | Mean | SD | N | Mean | SD | N | Mean | SD | N | Mean | SD |
| BODY: Body Length (cm) | 27 | 21.9 | 1.4 | 14 | 22.2 | 1.9 | 29 | 19.9 | 1.6 | 52 | 20.9 | 2.7 |
| FEMUR: Femur Length (cm) | 27 | 6.5 | 0.6 | 14 | 6.3 | 0.5 | 29 | 5.5 | 0.7 | 54 | 5.8 | 0.6 |
| FO: Fronto-Occipital Distance (mm) | 27 | 42.6 | 2.9 | 14 | 40.2 | 2.6 | 24 | 39.7 | 2.2 | 50 | 39.8 | 2.1 |
| FOOT: Foot Length (mm) | 24 | 61.1 | 5.6 | 14 | 55.4 | 3.2 | 27 | 54.7 | 4.6 | 54 | 54.2 | 3.7 |
| FOREARM: Distance Wrist to Elbow (cm) | 27 | 5.2 | 0.4 | 14 | 4.8 | 0.3 | 29 | 4.5 | 0.4 | 54 | 4.6 | 0.5 |
| HAND: Distance Wrist to Tip of Longest Finger (mm) | 21 | 30.5 | 15.1 | 13 | 36.3 | 2.7 | 28 | 35.4 | 3.3 | 48 | 35.2 | 4.0 |
| HUMERUS: Humerus Length (cm) | 26 | 5.4 | 0.8 | 14 | 5.3 | 0.3 | 29 | 4.7 | 0.5 | 54 | 4.6 | 0.7 |
| IC: Intercranial Distance (mm) | 27 | 33.1 | 1.3 | 14 | 30.2 | 1.9 | 29 | 27.4 | 2.2 | 54 | 28.4 | 1.6 |
| JAW: Width of the Jaw (mm) | 23 | 23.7 | 3.9 | 14 | 25.7 | 3.1 | 29 | 22.2 | 2.9 | 52 | 22.9 | 2.3 |
| TAIL: Tail Length (cm) | 26 | 32.3 | 1.7 | 13 | 30.7 | 3.2 | 24 | 27.4 | 2.9 | 51 | 27.7 | 3.2 |
| TIBIA: Tibia Length (cm) | 27 | 7.2 | 0.5 | 14 | 7.1 | 0.3 | 29 | 6.6 | 0.6 | 54 | 6.5 | 0.6 |
| WEIGHT: Weight (g) | 25 | 440.6 | 66.8 | 14 | 386.2 | 63.0 | 30 | 322.6 | 65.2 | 54 | 308.4 | 68.1 |
| ZYG: Maximal zygomatic breadth (mm) | 23 | 31.4 | 2.6 | 14 | 30.1 | 3.3 | 29 | 28.7 | 1.5 | 51 | 28.6 | 2.1 |

*C. jacchus* and *C. penicillata* trait means (Supplementary Table S5 and Fig. 4). Overall, *C. jacchus* and *C. penicillata* tend to be the smallest among all taxa across morphological traits (Table 2 and Fig. 4).

Among hybrid taxa, *C. aurita* hybrids tended to have the largest median and mean values for all measured traits (Fig. 4 and Table 3). On the other hand, *C. penicillata* x *C. jacchus* hybrids showed the smallest median values and mean for most traits (Fig. 4 and Table 4). For hybrids and their parental species, *C. aurita* hybrids were not significantly different from *C. aurita* nor *C. geoffroyi* for any trait means based on post-hoc tests (Fig. 4, and Supplementary Table S5). There was a significant post-hoc difference in WEIGHT and FEMUR means between *C. aurita* hybrids and *C. jacchus* Supplementary Table S5). A post-hoc difference in WEIGHT means was also significant between *C. aurita* hybrids and *C. penicillata* (Supplementary Table S5) For *C. geoffroyi* x *C. penicillata* hybrids and *C. geoffroyi*, there were no significant post-hoc differences for any trait means (Supplementary Table S5). On the other hand, *C. geoffroyi* x *C. penicillata* hybrids were significantly different from *C. penicillata* for almost half of measured traits (Supplementary Table S5). There were no significant differences between *C. jacchus* x *C. penicillata* hybrids and either of the parental species in post-host testing (Supplementary Table S5).

**Table 3.**
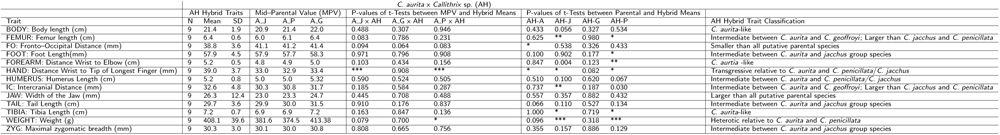
Summary of means, standard deviations (SD), sample numbers (N), mean mid-parental values (MPV) for thirteen morphological traits in *C. aurita* x *Callithrix* sp. hybrids (AH). In the “Mid-Parental Value (MPV)” columns “A J” is the MPV between *C. aurita* and *C. jacchus*, “A P” is the MPV between *C. aurita* and *C. penicillata*, and “A G” is the MPV between *C. aurita* and *C. geoffroyi*. In the “P-values of t-Tests between MPV and Hybrid Means” columns “AJ x AH” represents p-values from t-tests between AH hybrid trait means to the A J MPV, “A G x AH” represents p-values from t-tests between AH hybrid trait means to the A G MPV, and “A P x AH represents p-values from t-tests from AH hybrids to the AP MPV. In the “P-values of t-Tests between Parental and Hybrid Means” column “AH-A” indicates p-values of Welch’s t-tests between AH hybrids and *C. aurita* trait means. The “AH-J” column indicates p-values of Welch’s t-tests between AH hybrids and *C. jacchus* trait means, the “AH-G” column indicates p-values of Welch’s t-tests between AH hybrids and *C. geoffroyi*, the “AH-P” column indicates p-values of Welch’s t-tests between AH hybrids and *C. penicillata* trait means. Significant p-values are indicated as “*” for p-value*<*0.05, as “**” for p-value*<*0.01, and as “***” for p-value*<*0.001. “Cm” refers to centimeters, “mm” to millimeters, and “g” to grams.

**Table 4.**
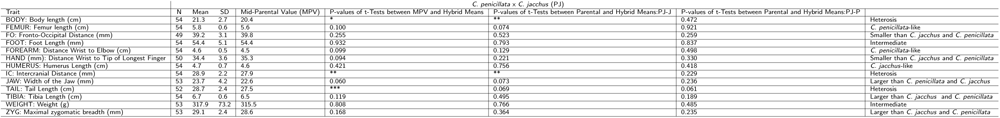
Summary of means, standard deviations (SD), sample numbers (N), mean mid-parental values (MPV) for thirteen morphological traits in *C. penicillata* x *C. jacchus* hybrids (PJ). The “P-values of t-Tests between MPV and Hybrid Means” column shows p-values from t-tests between PJ hybrids and the MPV. The “P-values of t-Tests between Parental and Hybrid Means:PJ-J” column represents Welch’s t-tests p-values between *C. jacchus* and PJ hybrids. The “P-values of t-Tests between Parental and Hybrid Means:PJ-P” column represents p-values for Welch’s t-tet between *C. penicillata* and PJ hybrids. Significant p-values are indicated as “*” for p-value*<*0.05, “**” for p-value*<*0.01, and “***” p-value*<*0.001. “Cm” refers to centimeters, “mm” to millimeters, and “g” to grams.

### Intermediacy, Heterosis, Dysgenesis, and Transgressive Segregation between Parental and Hybrid <u>Callithrix</u> Morphometric Traits (Aim 3)

Among *C. aurita* x *Callithrix* sp. hybrids (Table 3), we found evidence for transgressive segregation in the hand trait (HAND) when parental species combinations were either *C. aurita*/*C. penicillata* or *C. aurita*/*C. jacchus*. We also found evidence for heterosis in the WEIGHT trait if the parental species combination was *C. aurita-C. penicillata* (Table 3). The means of remaining traits for *C. aurita* x *Callithrix* sp. hybrids showed a tendency of being intermediate between all putative parental species or being larger than trait means of *C. jacchus* and *C. penicillata*. For *C. jacchus* x *C. penicillata* hybrids (Table 4), most trait means were larger than either of the parental species, though only a subset of these traits was significantly larger. Heterosis among these hybrids is shown in the TAIL, BODY, and IC traits, and no traits displayed evidence for dysgenesis. FOOT and WEIGHT traits were intermediate between *C. jacchus* x *C. penicillata* hybrids and their parental species. For *C. penicillata* x *C. geoffroyi* hybrids (Table 5), we found evidence for heterosis in the ZYG, TAIL, TIBIA, and FEMUR traits, while FO and JAW showed evidence of dysgenesis. The BODY, WEIGHT, and IC traits in *C. penicillata* x *C. geoffroyi* hybrids were intermediate, and none were transgressive.

**Table 5.**
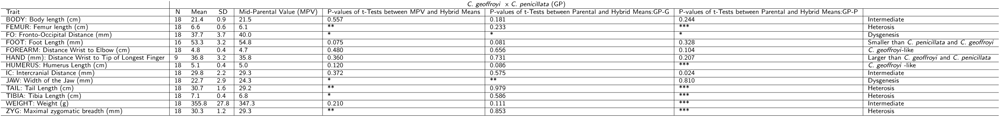
Summary of means, standard deviations (SD), sample numbers (N), mean mid-parental values (MPV) for thirteen morphological traits in *C. penicillata* x *C. geoffroyi* hybrids (GP). The “P-values of t-Tests between MPV and Hybrid Means” column shows p-values from t-tests between GP hybrids and the MPV. The “P-values of t-Tests between Parental and Hybrid Means:GP-G” column represents p-values of Welch’s t-tests between *C. geofforyi* and GP hybrids. The “P-values of t-Tests between Parental and Hybrid Means:GP-P” column represents p-values from Welch’s t-tests *C. geoffroyi* and GP hybrids. Significant p-values are indicated as “*” for p-value*<*0.05, “**” for p-value*<*0.01, and “***” p-value*<*0.001. “Cm” refers to centimeters, “mm” to millimeters, and “g” to grams.

The PCA plot *C. jacchus* x *C. penicillata* hybrids and their parental species as well as the positive loadings of PC1 (38.12% of variance) indicated a high degree of overlap between hybrids and parental species for overall size (Fig. 5A and Supplementary Table S6). The hybrids on average occupy an intermediate space shape between their parental species, but hybrid variation magnitude exceeds that of the parental species (Supplementary Table S7). Other PCs beyond PC1 of the *C. jacchus* x *C. penicillata* hybrids and parental species PCA combined positive and negative values indicating that they portray aspects of shape (Supplementary Table S7).

**Figure 5.**
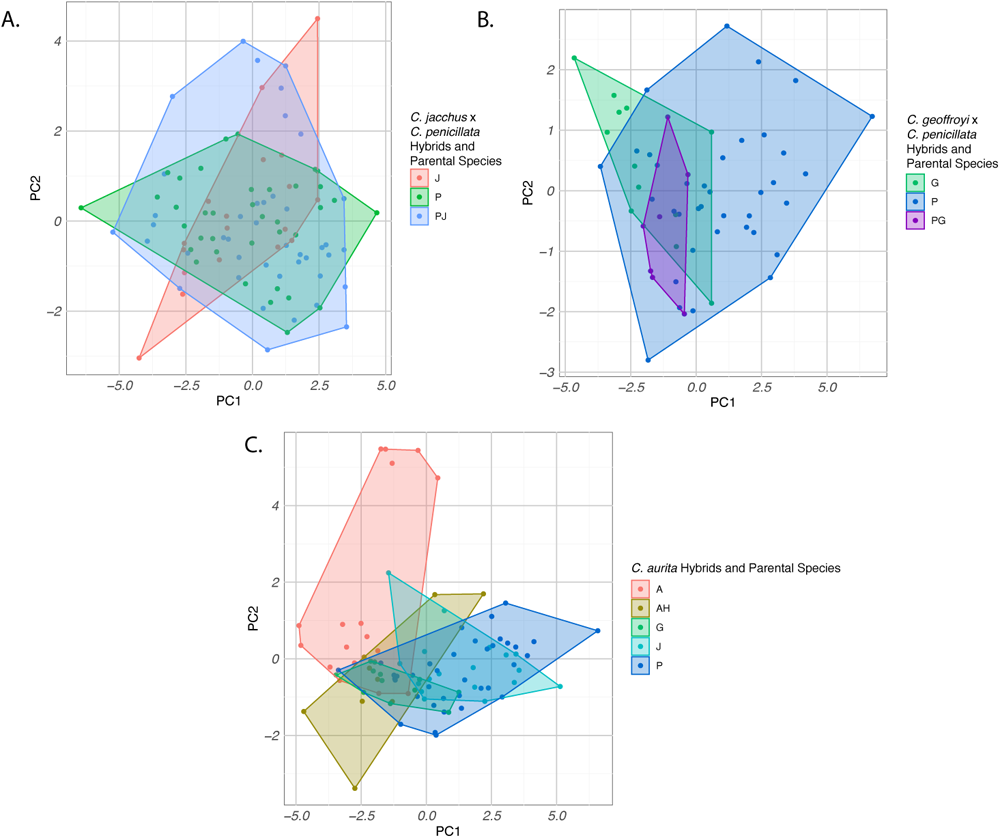
PCA plots for 13 morphological traits in Callithrix hybrids and their species. Bivariate plots of scores for the _rst two principal components factors are labelled and colored to indicate taxon a_liation. Plot A shows C. jacchus, C. penicillata and their hybrids. Plot B shows C. penicillata, C. geo_royi, and their hybrids. Plot C shows C. aurita, C. jacchus, C. geo_royi, C. penicillata, and their hybrids. Plot legends indicate taxon a_liation as follows: A= C. aurita, G= C. geo_royi, P= C. penicillata, JP= C. jacchus x C. penicillata hybrids, PG=C. penicillata x C. geo_royi hybrids, AH= C. aurita hybrids.

The PCA for *C. penicillata*, *C. geoffroyi*, and their hybrids (Fig. 5B) shows some separation between the two parental species along PC1 (42.90%), with larger *C. geoffroyi* towards the left and smaller *C. penicillata* towards the right. PCA eigen-values for this analysis are shown in Supplementary Table S8. Hybrids fall in between the two parental species along PC1 and PC2, indicating that the magnitude of variation in the sampled hybrids does not exceed that of parental species (Supplementary Table S8). The negative loadings of PC1 of this PCA may portray aspect of overall size. PC2 shows positive and negative values which may portray shape aspects among *C. penicillata*, *C. geoffroyi*, and their hybrids (Supplementary Table S9).

The PCA plot of the four study species and *C. aurita* x *Callithrix* sp. hybrids (Fig. 5C) shows most overlap between the three *jacchus* group species to the exclusion of *C. aurita* along PC1 (42.90% of variance). PC1 seems to be influenced by both size and shape of the marmosets (Fig. 5C). The hybrids cluster closest to *C. aurita* toward the left side. PCA eigenvalues for this analysis are shown in Supplementary Table S10. All negative loading on PC1 indicate that this may be an overall size component (Supplementary Table S11). PC2 (17.76% of variability) seems heavily influenced by jaw, FO, and hand (Supplementary Table S11). The magnitude of *Callithrix aurita* hybrid variation magnitude exceeds that of the all parental species (Fig. 5C).

### Callithrix Species Mitogenomic Genetic Distances (Aim 4)

Mean pairwise mitogenomic genetic distance between *C. jacchus*, *C. penicillata*, *C. geoffroyi*, and *C. aurita* are listed in Table 6. These measures show that *C. jacchus* and *C. penicillata* possessed the smallest mean distance out of all pairwise comparisons. Then *C. geoffroyi* had the same genetic distance from both *C. jacchus* and *C. penicillata*. Finally, *C. aurita* was the most genetically removed from all three other species.

**Table 6.** Species mean pairwise genetic distances of four *Callithrix* species based on previously published mitogenomic haplotypes which include a subset of marmosets sampled in this study.

|  | *C. aurita* | *C. geoffroyi* | *C. jacchus* | *C. penicillata* |
| --- | --- | --- | --- | --- |
| *C. aurita* |  |  |  |  |
| *C. geoffroyi* | 0.059 |  |  |  |
| *C. jacchus* | 0.060 | 0.018 |  |  |
| *C. penicillata* | 0.059 | 0.018 | 0.014 |  |

## Discussion

### Pelage Variation in Callithrix Species and Hybrids

*Callithrix* hybrids pelage patterns and coloration incorporate parental phenotypes into novel combinations [26, 28, 34, 35], but the functional consequences of this phenotypic variation are still unclear. Hypotheses that explain the function of phenotypic variation in primate coloration include protection, communication, and character displacement [45, 46, 47]. For example, Gloger’s rule predicts that endothermic animals, including primates, will be darker in wetter, more humid locations [45, 48], which may play a role in thermoregulation [49]. Among *Callithrix* marmosets, *Callithrix aurita* has the darkest overall pelage, and occurs in some of the highest average rainfall regions of natural *Callithrix* geographical ranges [26, 50]. On the other hand, *C. jacchus* and *C. penicillata*, which inhabit the semi-arid Caatinga and Cerrado biomes [26, 50], show lighter pelage than other *Callithrix* species. Additionally, *C. jacchus* and *C. penicillata* do indeed show lighter pelage around the eyes and darker tones around the mouth and nose, as expected for primates found in semi-arid regions [46]. As a portion of our sampled individuals came from captive settings or from unknown provenance, our current data set cannot be used for testing hypotheses of phenotypic variation in marmoset hybrids. However, a future study direction would be to develop statistical and/or artificial intelligence models to understand how environment (e.g. Gloger’s rule) and genetic variables influence phenotypic variation of pelage pattern and coloration inside and outside of marmoset hybrid zones. Under the character displacement, the intricacy of pelage coloration is used by individuals to distinguish conspecifics from heterospecifics to reduce the probability of hybridization [45]. Thus, possible directions for future studies include the integration of phenotypic data with measures of reproductive fitness and mate choice for marmoset hybrids and species at natural and anthropogenic hybrid zones. Such studies could shed light on whether specific marmoset phentoypic features are associated with reproductive success of hybrid and nonhybrid.

One study recently suggested that multigenerational marmoset hybrids experience a “greying out” of parental pelage coloration as hybridization goes on over time and that parental characteristics are only distinguishable in early generation hybrids [51]. However, data on pelage phenotypes presented in this study and previously published studies do not sustain this prediction. For example, in several late-generation natural and anthropogenic hybrid zones between *jacchus* group species, parental phenotype and genotype combinations, respectively, are uncoupled within hybrid populations and reshuffled into new combinations amongst hybrid individuals [26, 28, 34, 35]. Parental pelage characteristics and coloration are still observable in anthropogenic marmoset hybrid zones that have existed for over 30-40 years (that is about 45-60 marmoset generations assuming a marmoset generation time of 1.5 year), and that do not receive natural gene flow from parental species [34, 35]. The greyish marmoset hybrids exemplified by Vital et al. [51], are similar in pelage phenotype to the *C. aurita* x *jacchus* group hybrids we present in this study and also discussed in [28]. These marmosets hybrids are greyer in appearance than *jacchus* group hybrids, but also retain pelage characteristics indicative of ancestry from both *aurita* and *jacchus* group marmoset species. Genomic data on global admixture levels for the *C. aurita* x *Callithrix* sp. hybrids in our study (unpublished data, Malukiewicz) suggest that these are likely late generation hybrids, which goes against any progressive greying-out of pelage hypothesis in such hybrids.

### Morphometric Variation in Callithrix Species

Marmoset cranial shape and musculature, dentition, in addition to digestive features [52, 53, 54, 55], support *Callithrix* exudivory by allowing marmosets to gouge and scrape hard plant surfaces to access and digest natural exudate sources made of hard to digest oligosaccharides [52, 56, 57, 58, 55, 54, 53, 59, 60, 61, 62, 63, 64]. However, interspecific differences in marmoset cranial shape and dentition *Callithrix* species are linked to intersepcific differences in exudivory specializaiton [56, 65, 66, 67], with *C. jacchus* and *C. penicillata* representing the extreme of marmoset exudivory specialization and *C. aurita* being the least specialized [68]. *Callithrix penicillata* and *C. jacchus* have compressed braincases and more protruding dentition in comparison to *Callithrix aurita* and *C. flaviceps* [56]. Specifically in *C. jacchus*, the cranial musculoskeletal configuration allows for the use of extreme wide jaw gapes to gouge tree holes with the anterior dentition. In our results for cranial traits (IC, FO, ZYG, and JAW) [58, 60], we saw significant pairwise differences between *C. aurita C. jacchus* and *C. aurita C. penicillata* comparisons while all pairwise comparisons between *C. jacchus* and *C. penicillata* were not significant. Other studies have reported either no significant differences or a high degree of overlap in *C. jacchus* and *C. penicillata* cranial and dental traits and that these species are morphological distinct in such traits from *C. aurita* [66, 67, 56]. We attribute the differences seen in craniofacial morphology of marmoset species in our results to differences in exudivory specialisation between these species [27, 68].

Primate exudivores tend to be small in size [63], and in our study the most extreme marmoset exudivores, *C. jacchus* and *C. penicillata* were on average the smallest for all thirteen morphological traits. Then as with cranial traits, these two species were the only pair which did not possess any significant pairwise trait differences for post-cranial traits. On the other hand, *C. aurita* as the least specialized species for exudivory, tended on average to be the largest for most of the thirteen studied morphological traits. These species respectively represent the two relative extremes of exudivory in *Callithrix*, with the other marmoset species falling somewhere in between as far as exudate consumption [27]. Morphologically, *C. geoffroyi* fell in between the rest of the species included here. Other morphological studies of the marmoset cranium show that *C. flaviceps* is most similar to *C. aurita* and *C. kuhlii* is closer to the other four *Callithrix species* [56, 67, 67, 65]. These trends also reflect level of exudivory specialization in these other species [27].

### Morphometric Variation in Callithrix Hybrids and their Parental Species

Underlying differences in the degree of genetic similarity between parental taxa of hybrids are important factors in determining patterns of phenotypic variation in hybrids [10, 20, 21, 23]. Our results show that patterns of hybrid phenotypic variation relative to parental species are not consistent among marmoset hybrids with differing parental species ancestries. We see the least amount of MPV deviation in hybrids with the least mitogenomic genetic distance between the parental species, that being *C. jacchus* and *C. penicillata*, with several intermediate or parentalspecies-like traits and three traits with heterosis. Due to their genetic closeness and adaptive similarities, there is likely less breakdown of co-adaptive gene complexes between *C. jacchus* and *C. penicillata* than between other pairings of *Callithrix* parental species in our sample. We also probably see less heterosis in *C. jacchus* x *C. penicillata* hybrids than in other hybrid types in our sample as there may be a lesser amount of differentially fixed alleles between *C. jacchus* and *C. penicillata* than between other marmoset species.

In line with the expectation that larger differences in gene frequencies between parental populations contribute to the occurrence of heterosis and dysgenesis in hybrids [21], *C. penicillata* x *C. geoffroyi* hybrids, whose parental species possess larger mitogenomic distance that *C. jacchus-C. penicillata*, show four traits with heterosis, two with dysgenesis, and three intermediate traits. In the latter set of traits, three were closer to *C. geoffroyi* means than *C. penicillata* means. A previous study of *C. penicillata* x *C. geoffroyi* hybrids in the same sampling locality as ours also found that for traits which fell within the parental species range, hybrids were closer to *C. geoffroyi* than *C. penicillata* [35]. For the *C. aurita* hybrids, WEIGHT was heterotic in *C. aurita*-*C. penicillata* contrasts, which are putative parental species pairs with a relatively high level of genetic differentiation. Due to less genetic and adaptive similarity between *C. penicillata* and *C. geoffroyi* and *C. aurita* and all *jacchus* group species, respectively, relative to *C. jacchus* and *C. penicillata*, our results suggests some breakdown of co-adaptive gene complexes, and higher number of different alleles that have been fixed between the former than latter pair of parental species.

Transgression in hybrids is expected to increase with greater genetic distance between interbreeding parental species due to complementary gene action or epistasis [20]. We observed transgression in the HAND trait of *C. aurita* hybrids between *C. aurita*-*C. jacchus* and *C. aurita*-*C. penicillata* contrasts, which represent the most genetically distant pairing of parental species in our sample. PCA plots of *C. jacchus* and *C. penicillata* show that most hybrids fall within the range of parental species phenotypic variation, but a few extreme hybrid individuals outside of the parental range represent transgressive individuals. Interestingly, we did not see indication of trangressive hybrids in PCA plots of *C. geoffroyi* x *C. penicillata* hybrids and parental species, while Fuzessy et al. [35] did. This difference maybe due to a larger number of hybrids sampled by Fuzessy et al (N=40) than in this study (N=18). For *aurita* x *jacchus* group hybrids, most of these individuals are transgressive that fall outside the phenotyptic range of all four parental species. Thus, transgressive hybridization in marmosets, when considering morphometric shape and size in terms of genetic relatedness between parental species, follows theoretical expectations.

### Implications of Understanding Marmoset Hybrid Pelage and Morphometric Diversity

Our results based on *Callithrix* show that indeed expression of morphometric traits differs in hybrids resulting from interbreeding between different combinations of closely-related parental species that differ in genetic distance. Temporal divergence between parental marmoset species included in this study tracks positively with their level of genetic distance [27]. Further, experimental hybrid crosses showed that *C. jacchus* and *C. penicillata* hybridize relatively more easily than other *Callithrix* species pairing, and their hybrid progeny also show relatively less physical abnormalities (see [28]). Thus, our empirical data and past experimental data suggest that less developmental disturbances can be expected in hybrids of species that have diverged relatively more recently. Given the various anthropogenic hybrids found across southeastern Brazil, *Callithrix* marmosets represent a system where this question can be explored more directly for phenotypes related to anatomy and beyond experimental setting.

Marmoset morphological adaptations related to exudivory may be one set of traits which have important implications for viability and adaptability across different types of wild *Callithrix* hybrids in anthropogenic hybrid zones. Plant exudates are an important nutritional resource for natural populations *C. jacchus* and *C. penicillata* [69, 70, 71], and are also likely an important fallback food for exotic populations of these species in the southeastern Brazilian Atlantic Forest. Fallback foods are considered “nutritional resources for which a species has evolved specific masticatory and digestive adaptations, and are consumed principally when preferred foods are scarce” [72]. This study and another recently published work [36] show that cranial traits important for marmoset exudivory (e.g., zygomatic breath and width of jaw [59, 58, 60]) are largely not affected by heterosis, dysgenesis, or trangression in *C. jacchus* x *C. penicillata* hybrids. This pattern is in contrast to the relatively more frequent occurrence of heterosis we observed in the post-cranial traits of *C. jacchus* x *C. penicillata* hybrids. In the other types of hybrids included in this study, we observed relatively more heterosis, dysgenesis, and trangression in both cranial and post-cranial traits. Based on these patterns, the question arises if there is more selective pressure in *C. jacchus* x *C. penicillata* hybrids to minimize developmental disturbance of cranial morphology than post-cranial morphology. Retaining the exudivory specialization of their parental species likely enables *C. jacchus* x *C. penicillata* hybrids to use plant exudates as a fallback food, and contribute to these hybrids being the most common hybrid type present in anthropogenic *Callithrix* hybrid zones in the southeastern Brazilian Atlantic Forest [28, 27, 73, 36]. These factors may affect the ability of such hybrids to successfully exploit plant exudates as fallback foods relative to *C. jacchus* and *C. penicillata* hybrids. As a result, these other types of hybrids may be less adaptable to anthropogenic hybrid zones and in the longer run less viable.

Further tests of selective pressuring on cranial and post-cranial morphological traits in marmosets should combine phylogenetic, genomic, demographic, and phenotypic data from sampled hybrids and their parental species. Future studies should also consider underlying genetic architecture of a given trait, level of admixture, and the generational age of hybrids. Combining these factors will provide a fuller understanding of hybrid phenotypic expression, and provide insight into how natural animal populations may evolve as anthropogenic hybridization continues to increase. For marmosets themselves, establishing a firm understanding of phenotypic differences and variability in both *Callithrix* species and hybrids is important for both evolutionary, conservation, and applied reasons. Anthropogenic marmoset hybrids and exotic marmosets regularly fill up governmental and zoological captive facilities in Brazil and marmoset species such as *C. jacchus* are usually kept in biomedical facilities outside of Brazil. Pelage colors and patterns that are easily observable and distinguishable are usually the first key characteristics to classify a marmoset individual as either a hybrid or non-hybrid as well as the likely ancestry of that species. Anthropogenic hybrids pose ecological and conservation challenges, particularly in southeastern Brazil, but natural marmoset hybrids are also found along the entire geographical *Callithrix* range. Thus proper identification of marmoset hybrid and ancestral status is fundamental in execution of any marmoset conservation and population management plans in and out of captivity. Our suggestions to this end include adopting and developing quantitative approaches and tools towards identification and taxonomic classification of marmosets, as most approaches still depend on subjective, qualitative descriptions which are subject user error. A new direction we are currently involved in is the development of a machine-learning internet and phone app to help biological and clinical workers easily identify marmosets. Ideally, phenotypic data should be combined with mitochondrial and nuclear genome data in identification and classification of marmosets, as phenotypic data is not fully reliable to this end as cryptic hybridization does occur in marmosets [28, 74].

## Declarations

### Ethics approval and consent to participate

Animals were sampled under the approval of the Arizona State University Institutional Animal Care and Use Committee Animals (ASU IACUC, protocols #11-1150R, 15-144R), Brazilian Environmental Ministry (SISBIO protocols #47964-2 and #28075-2), and a CPRJ internal review. All methods were carried out in accordance with relevant guidelines and regulations. All methods are reported in accordance with ARRIVE guidelines (https://arriveguidelines.org) for the reporting of animal experiments.

### Consent for publication

Not applicable.

## Availability of data and materials

All data generated or analysed during this study are included in this published article [and its supplementary information files].

## Author’s contributions

JM, KW, IOS, and RRA formulated the idea for the study. KW and RRA formulated analytical aspects of this work. KW conducted data analyses of this work. JM collected samples, obtained funding, conducted analyses, and wrote the original manuscript. ILCA aided in field and data collection. NHAC provided field assistance and logistical support. FTD, LSF, HRF, LM, VSP, and BP provided veterinary oversight and logistical support for field collections at PET. CVM assisted in phenotypic identification of hybrids, gave field assistance, veterinary oversight, and aided in data collection. CSI gave access and provided logistical and veterinary support to collect samples from animals kept at Guarulhos Zoo. AP gave access to collect samples at Centro de Primatologia do Rio de Janeiro. SBM provided logistical support and veterinary assistance to collect samples from animals kept at the Centro de Primatologia do Rio de Janeiro. TZ, MSN and JLS provided access, logistical support, and veterinary assistance at DEPAVE. PAN and LCMP gave access and provided logistical support to collect samples from animals kept at CEMAFAUNA. DLS and MOMS provided assistance in field and data collections. JR contributed towards writing the original manuscript. AAQ provided veterinary oversight and collection assistance at CEMAFAUNA. VB and IOS provided field and logistical assistance. All authors read and approved the final manuscript.

## Supporting information

Supplementary_Figure_S1_legend.txt

Supplementary_Figure_S2_legend.txt

Supplementary Table S1.tsv

Supplementary Table S2.tsv

Supplementary Table S3.tsv

Supplementary Table S4.tsv

Supplementary Table S5.tsv

Supplementary Table S6.tsv

Supplementary Table S7.tsv

Supplementary Table S8.tsv

Supplementary Table S9.tsv

Supplementary Table S10.tsv

Supplementary Table S11.tsv

## Acknowledgements

We would like to thank the Callitrichid Research Center, the Golden Lion Tamarin Association, CEMAFAUNA, Rio de Janeiro Primatology Center, the Guarulhos Municipal Zoo, the Beagle Lab at UFV, and many biologists, field technicians, veterinarians, and other individuals that made this research possible. We would like to thank Adrielle M. Cezar for early comments on this work.

## Additional Files

Additional file 1 — Supplementary Figure S1.pdf

Pictures showing labeled facial regions used for phenotypic identification of sampled hybrids. *Callithrix* species were distinguished by: (1) color of the lateral sides of the face; (2) coloration in the frontal and back portions of the vertex; (3) coloration, shape, and volume of the auricular tufts; (4) presence/absence of a white forehead marking; (4) coloration of the orbital region; and (6) coloration of the menton region.

Additional file 2 — Supplementary Figure S2.pdf

Morphological variable normal QQ plots for thirteen morphological traits used in this study.

Additional file 3 — Supplementary Table S1.tsv

Table S1. Metadata and individual morphological trait measures for sampled marmosets. The ‘Individual’ column gives ID of each sampled individual. The ‘Place of Collection’ column indicates whether an individual was sampled in the wild, at a captive facility, or came from the wild and then was transferred to a captive facility. The Guarulhos Municipal Zoo is located in Guarulhos, São Paulo, Brazil; CPRJ (Centro de Primatologia do Rio de Janeiro) is located in Guapimirim, Rio de Janeiro, Brazil; CEMAFAUNA (Centro de Conservação e Manejo de Fauna da Caatinga) is located in Petrolina, Pernambuco; DEPAVE (Prefeitura Municipal de São Paulo, Secretaria Municipal do Verde e Meio Ambiente - DEPAVE (Divisão Técnica de Medicina Veterińaria e Manejo da Fauna Silvestre) is located in São Paulo, São Paulo, Brazil; PET (Parque Ecoĺogico do Tiete) is located in São Paulo, São Paulo; PARNASO (Parque Nacional Serra dos Órgãos) is located in Teresopolis, Rio de Janeiro, Brazil. SERCAS (Setor de Etologia aplicada à Reintrodução e Conservação de Animais Silvestres) is located in Campos dos Goytacazes, Rio de Janeiro, Brazil. The ‘City’ and ‘State’ columns indicated where each individual was sampled. Abbreviations for Brazilian states in the ‘State’ column are as follows: Espírito Santo (ES), Minas Gerais (MG), Pernambuco (PE), Rio de Janeiro (RJ), São Paulo (SP). The ‘Taxon’ column indicates whether the sampled individual possessed a species or hybrid phenotype. Taxon abbreviations in this column are as follows: ‘A’ is *C. aurita*, ‘G’ is *C. geoffroyi*, ‘J’ is *C. jacchus*, ‘P’ is *C. penicillata*, ‘AH’ is *C. aurita* hybrid , ‘PJ’ is *C. jacchus* x *C. penicillata* hybrid, ‘PG’ is *C. penicillata* x *C. geoffroyi* hybrid, and ‘CC’ is *Callithrix* sp. x *Callithrix* sp. hybrid. The ‘Sex’ column indicates the sex of the sampled individuals (F=Female, M=Male). The ‘Age’ column indicates the age of the sampled individual (A=Age). The rest of the columns show individual measures for thirteen morphological traits (NA=No data Available). Abbreviations in each trait column match those described in the methods. Traits with left and right measures have been averaged for the analyses described in the methodology section of the main text.

Additional file 4 – Supplementary Table S2.tsv

Supplementary Table S2. List of previously published mitogenome haplotypes used to calculate genetic distances between the four marmoset species included in this study.

Additional file 5 – Supplementary Table S3.tsv

Supplementary Table S3. Description of key facial features, facial regions, and post-cranial body parts that characterize *Callithrix* species and hybrids with at least one known parental species.

Additional file 6 – Supplementary Table S4.tsv

Supplementary Table S4. Results of univariate Welch’s ANOVA test for differences across all *Callithrix* taxa for 13 morphometric traits.

Additional file 7 – Supplementary Table S5.tsv

Supplementary Table S5. Games-Howell post-hoc pairwise tests after Welch’s ANOVA to determine which comparisons between *Callithrix* taxa for thirteen individual traits are significant. ‘Trait’ column names of traits follow that of Supplementary Table S1. ‘Group 1’ and ‘Group2’ indicate which two taxa are being compared and abbreviations follow Supplementary Table S1. ‘Estimate’ column refers to the mean difference between the groups being compared, ‘conf.low’ column refers to lower limit of the confidence interval for the mean difference, ‘conf.high’ column refers to higher limit of the confidence interval for the mean difference, ‘p.adj’ is the adjusted p-value using Turkey’s method, and ‘p.adj.signif’ column indicates the significance level of adjusted p-values with ‘ns’ meaning note significant.

Additional file 8 – Supplementary Table S6.tsv

Supplementary Table S6. Eigenvalues and variance of principle components (PCs) for *C. jacchus* and *C. penicillata* hybrids and parental species.

Additional file 9 – Supplementary Table S7.tsv

Supplementary Table S7. Loadings of principal components (PCs) fo *C. jacchus* x *C. penicillata* hybrids and parental species

Additional file 10 – Supplementary Table S8.tsv

Supplementary Table S8. Eigenvalues and variance of principle components (PCs) for *C. geoffroyi* and *C. penicillata* hybrids and parental species.

Additional file 11 – Supplementary Table S9.tsv

Supplementary Table S9. Loadings of principal components (PCs) for *C. geoffroyi* x *C. penicillata* hybrids and parental species

Additional file 12 – Supplementary Table S10.tsv

Supplementary Table S10. Eigenvalues and variance of PCs (principle components) for *C. aurita*, *C. jacchus*, *C. geoffroyi* and *C. penicillata* hybrids and parental species.

Additional file 13 – Supplementary Table S11.tsv

Supplementary Table S11. Loadings of PCs for *C. geoffroyi*, *C. penicillata*, *C. jacchus*, and *C. aurita* hybrids and parental species.

Additional file 14 – Supplementary Figure S1 legend.txt

Figure legend for Supplementary Figure S1.

Additional file 15 – Supplementary Figure S2 legend.txt

Figure legend for Supplementary Figure S2.

Additional file 16 – morphometricsv3 code.Rmd

Code of R analyses described in this work.

## References

1. Mallet, J.: Hybridization as an invasion of the genome. Trends in Ecology & Evolution 20(5), 229–237 (2005). doi:10.1016/j.tree.2005.02.010

2. Crispo, E., Moore, J.-S., Lee-Yaw, J.A., Gray, S.M., Haller, B.C.: Broken barriers: Human-induced changes to gene flow and introgression in animals. BioEssays 33(7), 508–518 (2011). doi:10.1002/bies.201000154

3. Grabenstein, K.C., Taylor, S.A.: Breaking barriers: causes, consequences, and experimental utility of human-mediated hybridization. Trends in Ecology & Evolution 33(3), 198–212 (2018). doi:10.1016/j.tree.2017.12.008

4. McFarlane, S.E., Pemberton, J.M.: Detecting the true extent of introgression during anthropogenic hybridization. Trends in Ecology & Evolution 34(4), 315–326 (2019). doi:10.1016/j.tree.2018.12.013

5. Adavoudi, R., Pilot, M.: Consequences of hybridization in mammals: A systematic review. Genes 13(1), 50 (2021). doi:10.3390/genes13010050

6. Ottenburghs, J.: The genic view of hybridization in the anthropocene. Evolutionary Applications (2021). doi:10.1111/eva.13223

7. Moran, B.M., Payne, C., Langdon, Q., Powell, D.L., Brandvain, Y., Schumer, M.: The genomic consequences of hybridization. eLife 10 (2021). doi:10.7554/elife.69016

8. Kirkpatrick, M., Barton, N.H.: Evolution of a species’ range. The American Naturalist 150(1), 1–23 (1997). doi:10.1086/286054. PMID: 18811273.

9. Burgarella, C., Barnaud, A., Kane, N.A., Jankowsky, F., Scarcelli, N., Billot, C., Vigouroux, Y., Berthouly-Salazar, C.: Adaptive introgression: an untapped evolutionary mechanism for crop adaptation (2018). doi:10.1101/379966

10. Ackermann, R.R.: Phenotypic traits of primate hybrids: Recognizing admixture in the fossil record. Evolutionary Anthropology: Issues, News, and Reviews 19(6), 258–270 (2010). doi:10.1002/evan.20288

11. Warren, K.A., Ritzman, T.B., Humphreys, R.A., Percival, C.J., Hallgŕımsson, B., Ackermann, R.R.: Craniomandibular form and body size variation of first generation mouse hybrids: A model for hominin hybridization. Journal of Human Evolution 116, 57–74 (2018). doi:10.1016/j.jhevol.2017.12.002

12. Harvati, K., Ackermann, R.R.: Merging morphological and genetic evidence to assess hybridization in western eurasian late pleistocene hominins. Nature Ecology & Evolution 6(10), 1573–1585 (2022). doi:10.1038/s41559-022-01875-z

13. Kohn, L.A.P., Langton, L.B., Cheverud, J.M.: Subspecific genetic differences in the saddle-back tamarin (*Saguinus fuscicolli* s) postcranial skeleton. American Journal of Primatology 54(1), 41–56 (2001). doi:10.1002/ajp.1011

14. Bicca-Marques, J.C., Prates, H.M., de Aguiar, F.R.C., Jones, C.B.: Survey of *Alouatta caraya*, the black-and-gold howler monkey, and *Alouatta guariba clamitans*, the brown howler monkey, in a contact zone, state of rio grande do sul, brazil: evidence for hybridization. Primates 49(4), 246–252 (2008). doi:10.1007/s10329-008-0091-4

15. Kelaita, M.A., Cortés-Ortiz, L.: Morphological variation of genetically confirmed *Alouatta pigra*×*A. palliata* hybrids from a natural hybrid zone in tabasco, mexico. American Journal of Physical Anthropology 150(2), 223–234 (2012). doi:10.1002/ajpa.22196

16. Coğalniceanu, D., Stanescu, F., Arntzen, J.W.: Testing the hybrid superiority hypothesis in crested and marbled newts. Journal of Zoological Systematics and Evolutionary Research 58(1), 275–283 (2019). doi:10.1111/jzs.12322

17. Nikolakis, Z.L., Schield, D.R., Westfall, A.K., Perry, B.W., Ivey, K.N., Orton, R.W., Hales, N.R., Adams, R.H., Meik, J.M., Parker, J.M., Smith, C.F., Gompert, Z., Mackessy, S.P., Castoe, T.A.: Evidence that genomic incompatibilities and other multilocus processes impact hybrid fitness in a rattlesnake hybrid zone. Evolution (2022). doi:10.1111/evo.14612

18. Majtyka, T., Borczyk, B., Ogielska, M., Stöck, M.: Morphometry of two cryptic tree frog species at their hybrid zone reveals neither intermediate nor transgressive morphotypes. Ecology and Evolution 12(1) (2022). doi:10.1002/ece3.8527

19. Boel, C., Curnoe, D., Hamada, Y.: Craniofacial shape and nonmetric trait variation in hybrids of the japanese macaque (*Macaca fuscata*) and the taiwanese macaque (*Macaca cyclopis*). International Journal of Primatology 40(2), 214–243 (2019). doi:10.1007/s10764-019-00081-2

20. Stelkens, R.B., Schmid, C., Selz, O., Seehausen, O.: Phenotypic novelty in experimental hybrids is predicted by the genetic distance between species of cichlid fish. BMC Evolutionary Biology 9(1), 283 (2009). doi:10.1186/1471-2148-9-283

21. Cheverud, J.M., Jacobs, S.C., Moore, A.J.: Genetic differences among subspecies of the saddle-back tamarin (*Saguinus fuscicollis*):evidence from hybrids. American Journal of Primatology 31(1), 23–39 (1993). doi:10.1002/ajp.1350310104

22. BELL, M., TRAVIS, M.: Hybridization, transgressive segregation, genetic covariation, and adaptive radiation. Trends in Ecology & Evolution 20(7), 358–361 (2005). doi:10.1016/j.tree.2005.04.021

23. Rieseberg, L.H., Archer, M.A., Wayne, R.K.: Transgressive segregation, adaptation and speciation. Heredity 83(4), 363–372 (1999). doi:10.1038/sj.hdy.6886170

24. Rieseberg, L.H., Widmer, A., Arntz, A.M., Burke, B.: The genetic architecture necessary for transgressive segregation is common in both natural and domesticated populations. Philosophical Transactions of the Royal Society of London. Series B: Biological Sciences 358(1434), 1141–1147 (2003). doi:10.1098/rstb.2003.1283

25. Cortes-Ortiz, L., Duda, T.F., Canales-Espinosa, D., Garcia-Orduna, F., Rodriguez-Luna, E., Bermingham, E.: Hybridization in large-bodied new world primates. Genetics 176(4), 2421–2425 (2007). doi:10.1534/genetics.107.074278

26. Malukiewicz, J., Cartwright, R.A., Curi, N.H.A., Dergam, J.A., Igayara, C.S., Moreira, S.B., Molina, C.V., Nicola, P.A., Noll, A., Passamani, M., Pereira, L.C.M., Pissinatti, A., Ruiz-Miranda, C.R., Silva, D.L., Stone, A.C., Zinner, D., Roos, C.: Mitogenomic phylogeny of *Callithrix* with special focus on human transferred taxa. BMC Genomics 22(1) (2021). doi:10.1186/s12864-021-07533-1

27. Malukiewicz, J., Boere, V., de Oliveira, M.A.B., D’arc, M., Ferreira, J.V.A., French, J., Housman, G., de Souza, C.I., Jerusalinsky, L., de Melo, F.R., Valeņca-Montenegro, M.M., Moreira, S.B., de Oliveira e Silva, I., Pacheco, F.S., Rogers, J., Pissinatti, A., del Rosario, R.C.H., Ross, C., Ruiz-Miranda, C.R., Pereira, L.C.M., Schiel, N., de Fátima Rodrigues da Silva, F., Souto, A., Šlipogor, V., Tardif, S.: An introduction to the *Callithrix* genus and overview of recent advances in marmoset research. ILAR Journal 61(2-3), 110–138 (2020). doi:10.1093/ilar/ilab027

28. Malukiewicz, J.: A review of experimental, natural, and anthropogenic hybridization in *Callithrix* marmosets. International Journal of Primatology 40(1), 72–98 (2018). doi:10.1007/s10764-018-0068-0

29. Hershkovitz, P.: Comments on the taxonomy of brazilian marmosets (*Callithrix*, callitrichidae). Folia Primatologica 24(2-3), 137–172 (1975). doi:10.1159/000155687

30. Hershkovitz, P.: Living New World Monkeys (Platyrrhini). University of Chicago Press, Chicago (1977)

31. Passamani, M., Aguiar, L., Machado, R., Figueiredo, E.: Hybridization between *Callithrix geoffroyi* and *Callithrix penicillata* in southeastern minas gerais, brazil. Neotropical Primates 5, 9–10 (1997)

32. Mendes, S.: Padroes biogeograficos e vocais em *Callithrix* do grupo *jacchus* (primates, callitrichidae)). PhD thesis, Dissertation. Universidade Estadual de Campinas (UNICAMP) (1997)

33. Ruiz-Miranda, C.R., Affonso, A.G., de Morais, M.M., Verona, C.E., Martins, A., Beck, B.B.: Behavioral and ecological interactions between reintroduced golden lion tamarins (Leontopithecus rosalia linnaeus, 1766) and introduced marmosets (*Callithrix* spp, linnaeus, 1758) in brazil’s atlantic coast forest fragments. Brazilian Archives of Biology and Technology **49**, 99–109 (2006)

34. Malukiewicz, J., Boere, V., Fuzessy, L.F., Grativol, A.D., French, J.A., de Oliveira e Silva, I., Pereira, L.C.M., Ruiz-Miranda, C.R., Valeņca, Y.M., Stone, A.C.: Hybridization effects and genetic diversity of the common and black-tufted marmoset (*Callithrix jacchus* and *Callithrix penicillata*) mitochondrial control region. American Journal of Physical Anthropology 155(4), 522–536 (2014). doi:10.1002/ajpa.22605

35. Fuzessy, L.F., de Oliveira Silva, I., Malukiewicz, J., Silva, F.F.R., do Carmo P^onzio, M., Boere, V., Ackermann, R.R.: Morphological variation in wild marmosets (Callithrix penicillata and C. geoffroyi) and their hybrids. Evolutionary Biology 41(3), 480–493 (2014). doi:10.1007/s11692-014-9284-5

36. Cezar, A.M., Lopes, G.S., Cardim, S.S., Bueno, C., Weksler, M., Oliveira, J.A.: Morphological and genetic variation among callithrix hybrids in rio de janeiro, brazil. Evolutionary Biology 50(3), 365–380 (2023). doi:10.1007/s11692-023-09610-7

37. Yamamoto, M.: From dependence to sexual maturity: The behavioural ontogeny of callitrichidae. In: Rylands, A. (ed.) Marmosets and Tamarins: Systematics, Ecology and Behaviour, pp. 235–254. Oxford University Press, Oxford (1993)

38. Carvalho, R.: Conservação do saguis-da-serra-escuro (*Callithrix aurita* (primates)) – analise molecular e colormetrica de populaçoes do ĝenero callithrix e seus híbridos. dissertation, Universidade do Estado do Rio de Janeiro (2015)

39. Nagorsen, D.W., Peterson, R.L.: Mammal Collectors’ Manual : a Guide for Collecting, Documenting, and Preparing Mammal Specimens for Scientific Research. Royal Ontario Museum, (1980). Royal Ontario Museum

40. R Core Team: R: A Language and Environment for Statistical Computing. R Foundation for Statistical Computing, Vienna, Austria (2020). R Foundation for Statistical Computing. https://www.R-project.org/

41. Lääaräa, E.: Statistics: Reasoning on uncertainty, and the insignificance of testing null. Annales Zoologici Fennici 46(2), 138–157 (2009). doi:10.5735/086.046.0206

42. Kassambara, A.: Rstatix: Pipe-friendly Framework for Basic Statistical Tests. (2021). R package version 0.7.0. https://CRAN.R-project.org/package=rstatix

43. Tamura, K., Stecher, G., Kumar, S.: MEGA11: Molecular evolutionary genetics analysis version 11. Molecular Biology and Evolution 38(7), 3022–3027 (2021). doi:10.1093/molbev/msab120

44. Stecher, G., Tamura, K., Kumar, S.: Molecular evolutionary genetics analysis (MEGA) for macOS. Molecular Biology and Evolution 37(4), 1237–1239 (2020). doi:10.1093/molbev/msz312

45. Caro, T., Brockelsby, K., Ferrari, A., Koneru, M., Ono, K., Touche, E., Stankowich, T.: The evolution of primate coloration revisited. Behavioral Ecology 32(4), 555–567 (2021). doi:10.1093/beheco/arab029. https://academic.oup.com/beheco/article-pdf/32/4/555/39805493/arab029.pdf

46. Santana, S.E., Lynch Alfaro, J., Alfaro, M.E.: Adaptive evolution of facial colour patterns in neotropical primates. Proceedings of the Royal Society B: Biological Sciences 279(1736), 2204–2211 (2012). doi:10.1098/rspb.2011.2326.

47. Winters, S., Allen, W.L., Higham, J.P.: The structure of species discrimination signals across a primate radiation. eLife 9, 47428 (2020). doi:10.7554/eLife.47428

48. Delhey, K.: A review of gloger’s rule, an ecogeographical rule of colour: definitions, interpretations and evidence. Biological Reviews 94(4), 1294–1316 (2019). doi:10.1111/brv.12503.

49. Kamilar, J.M., Bradley, B.J.: Interspecific variation in primate coat colour supports Gloger’s rule. Journal of Biogeography 38(12), 2270–2277 (2011). doi:10.1111/j.1365-2699.2011.02587.x. eprint:. Accessed 2022-08-10

51. Alvares, C.A., Stape, J.L., Sentelhas, P.C., de Moraes Goņcalves, J.L., Sparovek, G.: Köppen’s climate classification map for Brazil. Meteorologische Zeitschrift, 711–728 (2013). doi:10.1127/0941-2948/2013/0507. Publisher: Schweizerbart’sche Verlagsbuchhandlung. Accessed 2022-08-10

51. Vital, O.V., Massardi, N.T., Brasileiro, S.L.S., Ĉorrea, T.C.V., Gjorup, D.F., Jerusalinsky, L., de Melo, F.R.: New records for *Callithrix aurita* and *Callithrix* hybriods in the region of viçosa, minas gerais, brazil. Neotropical Primates 26(2), 104–109 (2020)

52. Malukiewicz, J., Cartwright, R.A., Dergam, J.A., Igayara, C.S., Kessler, S.E., Moreira, S.B., Nash, L.T., Nicola, P.A., Pereira, L.C.M., Pissinatti, A., Ruiz-Miranda, C.R., Ozga, A.T., Quirino, A.A., Roos, C., Silva, D.L., Stone, A.C., Grativol, A.D.: The gut microbiome of exudivorous marmosets in the wild and captivity. Scientific Reports 12(1) (2022). doi:10.1038/s41598-022-08797-7

53. Power, M.L., Oftedal, O.T.: Differences among captive callitrichids in the digestive responses to dietary gum. American Journal of Primatology 40(2), 131–144 (1996). doi:10.1002/(sici)1098-2345(1996)40:2¡131::aid-ajp2¿3.0.co;2-z

54. Power, M.L., Myers, E.W.: Digestion in the common marmoset (*Callithrix jacchus*), a gummivore-frugivore. American Journal of Primatology 71(12), 957–963 (2009). doi:10.1002/ajp.20737

55. Caton, J.M., Hill, D.M., Hume, I.D., Crook, G.A.: The digestive strategy of the common marmoset, *Callithrix jacchus*. Comparative Biochemistry and Physiology Part A: Physiology 114(1), 1–8 (1996). doi:10.1016/0300-9629(95)02013-6

56. de Souza, V.B.: Variação do cr^anio e da mandíbula em *Callithrix* erxleben, 1777 (platyrrhini, callitrichidae): resultados de uma abordagem através de morfometria geométrica. Master’s thesis, Universidade Federal de Viçosa (2016)

57. Forsythe, E.C., Ford, S.M.: Craniofacial adaptations to tree-gouging among marmosets. The Anatomical Record: Advances in Integrative Anatomy and Evolutionary Biology 294(12), 2131–2139 (2011). doi:10.1002/ar.21500

58. Eng, C.M., Ward, S.R., Vinyard, C.J., Taylor, A.B.: The morphology of the masticatory apparatus facilitates muscle force production at wide jaw gapes in tree-gouging common marmosets (*Callithrix jacchus*). Journal of Experimental Biology 212(24), 4040–4055 (2009). doi:10.1242/jeb.029983

59. Taylor, A.B., Vinyard, C.J.: Comparative analysis of masseter fiber architecture in tree-gouging (*Callithrix jacchus*) and nongouging (*Saguinus oedipus*) callitrichids. Journal of Morphology 261(3), 276–285 (2004). doi:10.1002/jmor.10249

60. Vinyard, C.J., Wall, C.E., Williams, S.H., Mork, A.L., Armfield, B.A., de Oliveira Melo, L.C., Valeņca-Montenegro, M.M., Valle, Y.B.M., de Oliveira, M.A.B., Lucas, P.W., Schmitt, D., Taylor, A.B., Hylander, W.L.: The evolutionary morphology of tree gouging in marmosets. In: The Smallest Anthropoids, pp. 395–409. Springer, ??? (2009). doi:10.1007/978-1-4419-0293-120.

61. Pineda-Munoz, S., Alroy, J.: Dietary characterization of terrestrial mammals. Proceedings of the Royal Society B: Biological Sciences 281(1789), 20141173 (2014). doi:10.1098/rspb.2014.1173

62. CABANA, F., DIERENFELD, E.S., Wirdateti, DONATI, G., NEKARIS, K.A.I.: Exploiting a readily available but hard to digest resource: A review of exudativorous mammals identified thus far and how they cope in captivity. Integrative Zoology 13(1), 94–111 (2018). doi:10.1111/1749-4877.12264

63. Nash, L.T.: Dietary, behavioral, and morphological aspects of gummivory in primates. American Journal of Physical Anthropology 29(S7), 113–137 (1986). doi:10.1002/ajpa.1330290505

64. Smith, A.C.: Exudativory in primates: interspecific patterns. In: The Evolution of Exudativory in Primates, pp. 45–87. Springer, ??? (2010). doi:10.1007/978-1-4419-6661-2 3

65. Marroig, G., Cropp, S., Cheverud, J.M.: Systematics and evolution of the *jacchus* group of marmosets (platyrrhini). American Journal of Physical Anthropology 123(1), 11–22 (2003). doi:10.1002/ajpa.10146

66. Natori, M., Shigehara, N.: Interspecific differences in lower dentition among eastern-brazilian marmosets. Journal of Mammalogy 73(3), 668–671 (1992). doi:10.2307/1382041

67. Natori, M.: Craniometrical variations among eastern brazilian marmosets and their systematic relationships. Primates 35(2), 167–176 (1994). doi:10.1007/bf02382052

68. Rylands, A., Faria, D.: In: Rylands, A. (ed.) Habitats, feeding ecology, and home ranges size in the genus *Callithrix*, pp. 262–272. Oxford University Press, ??? (1993)

69. Bouchardet da Fonseca, G.A., Lacher, T.E.: Exudate-feeding bycallithrix jacchus penicillata in semideciduous woodland (cerradão) in central brazil. Primates 25(4), 441–449 (1984). doi:10.1007/bf02381667

70. Lamoglia, J.M., Boere, V., Picoli, E.A.d.T., de Oliveira, J.A., Silva Neto, C.d.M.e., Silva, I.d.O.: Tree species and morphology of holes caused by black-tufted marmosets to obtain exudates: Some implications for the exudativory. Animals 12(19), 2578 (2022). doi:10.3390/ani12192578

71. Pinheiro, H.L.N., Mendes Pontes, A.R.: Home range, diet, and activity patterns of common marmosets (callithrix jacchus) in very small and isolated fragments of the atlantic forest of northeastern brazil. International Journal of Ecology 2015, 1–13 (2015). doi:10.1155/2015/685816

72. Porter, L.M., Garber, P.A., Nacimento, E.: Exudates as a fallback food for callimico goeldii. American Journal of Primatology 71(2), 120–129 (2008). doi:10.1002/ajp.20630

73. Cezar, A.M., Pessoa, L.M., Bonvicino, C.R.: Morphological and genetic diversity in *Callithrix* hybrids in an anthropogenic area in southeastern brazil (primates: Cebidae: Callitrichinae). Zoologia 34, 1–9 (2017). doi:10.3897/zoologia.34.e14881

74. Malukiewicz, J., Cartwright, R.A., Dergam, J.A., Igayara, C.S., Nicola, P.A., Pereira, L.M.C., Ruiz-Miranda, C.R., Stone, A.C., Silva, D.L., Silva, F.d.F.R.d., Varsani, A., Walter, L., Wilson, M.A., Zinner, D., Roos, C.: Genomic skimming and nanopore sequencing uncover cryptic hybridization in one of world’s most threatened primates. Scientific Reports 11(1), 17279 (2021). doi:10.1038/s41598-021-96404-6. Number: 1 Publisher: Nature Publishing Group. Accessed 2022-08-10

