## Supplementary figures and images for "Pelage Variation and Morphometrics of Closely Related *Callithrix* Marmoset Species and Their Hybrids"

### Supplementary_Figure_S1.pdf

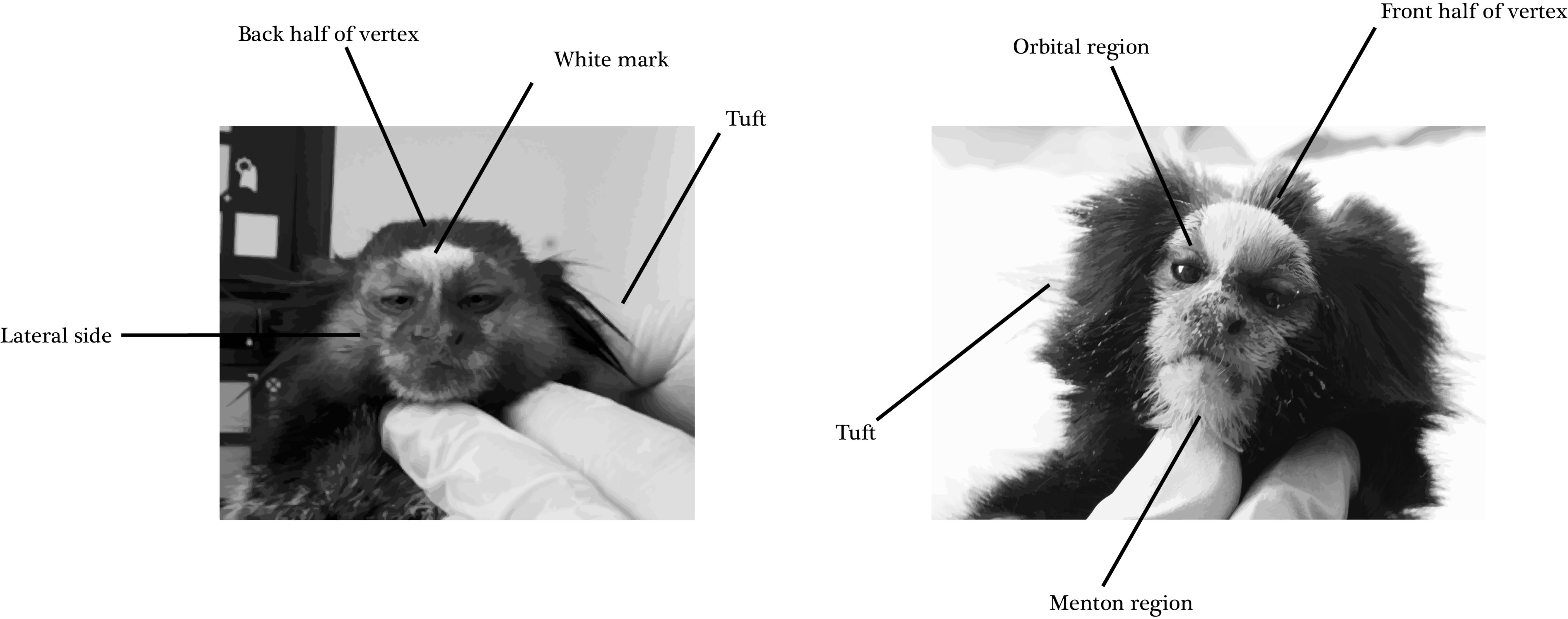

### Supplementary_Figure_S2.pdf

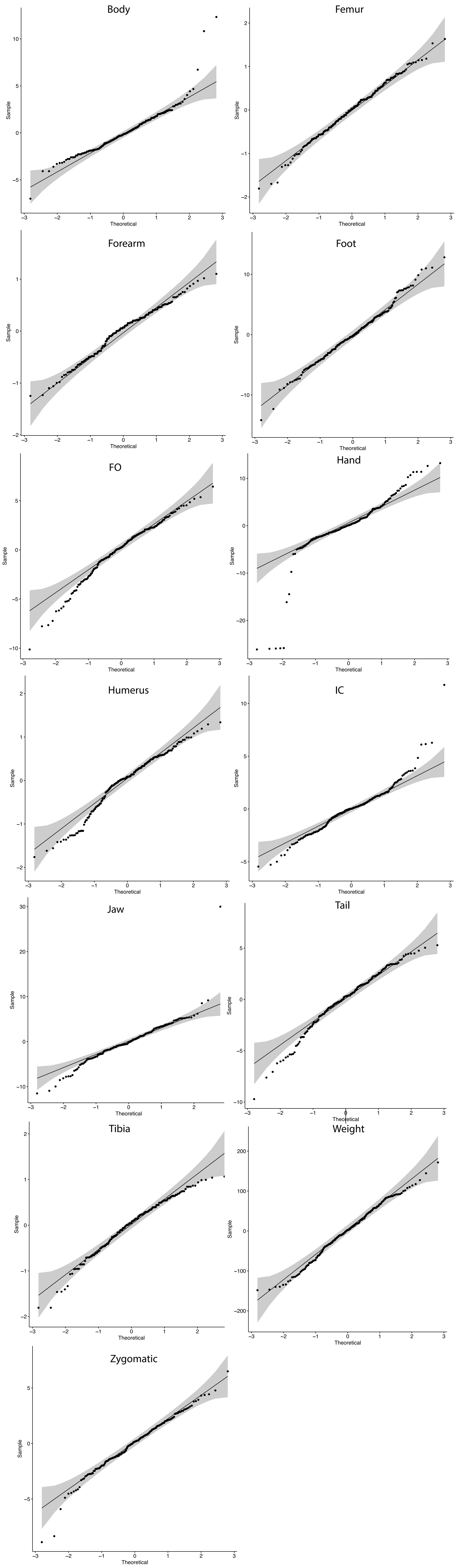
